# HNRNPA1 promotes recognition of splice site decoys by U2AF2 *in vivo*

**DOI:** 10.1101/175901

**Authors:** Jonathan M. Howard, Hai Lin, Garam Kim, Jolene M Draper, Maximilian Haeussler, Sol Katzman, Masoud Toloue, Yunlong Liu, Jeremy R. Sanford

**Affiliations:** Department of Molecular, Cellular and Developmental Biology, University of California Santa Cruz, 1156 High Street, Santa Cruz CA 95064; Department of Medical and Molecular Genetics, Indiana University School of Medicine, Indianapolis, IN 46202, USA; Center for Biomolecular Science and Engineering, UC Santa Cruz 1156 High Street, Santa Cruz CA 95064; Bioo Scientific Corporation, 7500 Burleston Rd, Austin, TX, 78744

**Keywords:** Alternative splicing, splice site recognition, HNRNPA1, U2AF

## Abstract

Alternative pre-mRNA splicing plays a major role in expanding the transcript output of human genes. This process is regulated, in part, by the interplay of *trans*-acting RNA binding proteins (RBPs) with myriad *cis*-regulatory elements scattered throughout pre-mRNAs. These molecular recognition events are critical for defining the protein coding sequences (exons) within pre-mRNAs and directing spliceosome assembly on non-coding regions (introns). One of the earliest events in this process is recognition of the 3’ splice site by U2 small nuclear RNA auxiliary factor 2 (U2AF2). Splicing regulators, such as the heterogeneous nuclear ribonucleoprotein A1 (HNRNPA1), influence spliceosome assembly both *in vitro* and *in vivo*, but their mechanisms of action remain poorly described on a global scale. HNRNPA1 also promotes proof reading of 3’ss sequences though a direct interaction with the U2AF heterodimer. To determine how HNRNPA1 regulates U2AF-RNA interactions in vivo, we analyzed U2AF2 RNA binding specificity using individual-nucleotide resolution crosslinking immunoprecipitation (iCLIP) in control- and HNRNPA1 over-expression cells. We observed changes in the distribution of U2AF2 crosslinking sites relative to the 3’ splice sites of alternative cassette exons but not constitutive exons upon HNRNPA1 over-expression. A subset of these events shows a concomitant increase of U2AF2 crosslinking at distal intronic regions, suggesting a shift of U2AF2 to “decoy” binding sites. Of the many non-canonical U2AF2 binding sites, Alu-derived RNA sequences represented one of the most abundant classes of HNRNPA1-dependent decoys. Splicing reporter assays demonstrated that mutation of U2AF2 decoy sites inhibited HNRNPA1-dependent exon skipping *in vivo*. We propose that HNRNPA1 regulates exon definition by modulating the interaction of U2AF2 with decoy or *bona fide* 3’ splice sites.

## Introduction

Precursor messenger RNA (pre-mRNA) splicing is catalyzed by a large macromolecular complex composed of five uridine-rich small nuclear ribonucleoprotein particles (U snRNPs) and myriad protein factors (Wahl et al. 2009). This process is required to excise intervening sequences (introns) from the pre-mRNA and ligate protein-coding sequences (exons). The 5’ and 3’ ends of introns are defined by nearly invariant GU and AG dinucleotides as well as the branch point sequence, respectively. Combinatorial protein-RNA interactions play important roles in defining the splice sites and branch point during spliceosome assembly. The 5’ splice site (GU) is recognized by serine and arginine-rich splicing factors (SR Proteins) and through base pairing interactions with the U1 snRNP (Eperon et al. 1993; Jamison et al. 1995). Similarly, the 3’ splice site is decoded by a combination of SR proteins and the U2 snRNP auxiliary factor (U2AF). After early (E) complex assembly, the branch point sequencing is specified through the interaction of U2AF, Splicing Factor 1/Branchpoint Binding Protein (SF1/BBP) and via base pairing with the U2 snRNP (Ruskin et al. 1988; Berglund et al. 1997). The responsibility for 3’ splice site recognition is shared by the two subunits of U2AF (Merendino et al. 1999; Wu et al. 1999; Zorio and Blumenthal 1999). The small and large subunit of U2AF, encoded by *U2AF1* and *U2AF2*, recognize the AG dinucleotide and the upstream polypyrimidine tract, respectively (Merendino et al. 1999; Wu et al. 1999; Zorio and Blumenthal 1999). These early steps in spliceosome assembly play critical roles in defining exon-intron boundaries.

HNRNPA1 is a well-characterized regulator of alternative splicing. One of the primary functions of HNRNPA1 is to prevent inclusion of specific exons (Mayeda and Krainer 1992; Caceres et al. 1994; Yang et al. 1994). Several models for HNRNPA1-dependent exon skipping have been described in the literature. For example, HNRNPA1 can compete with RBPs for binding to juxtaposed regulatory elements which would be normally occupied by splicing enhancer proteins such as SRSF1 (Eperon et al. 2000; Zahler et al. 2004). Another potential mechanism involves oligomerization and spreading of HNRNPA1 from an exonic splicing silencer across a regulated exon, thus antagonizing binding of splicing enhancers and spliceosomal factors, such as U2AF2 (Zhu et al. 2001; Okunola and Krainer 2009). Finally, HNRNPA1 binding to intronic splicing silencers on either side of an alternative exon can dimerize, causing “looping out” of an exon from the pre-mRNA and promote its exclusion from the final mature transcript (Blanchette and Chabot 1999). A related alternative splicing factor, PTBP1 also promotes exon skipping via a looping mechanism (Chou et al. 2000; Lamichhane et al. 2010). Perhaps most relevant to this work, HNRNPA1 promotes proofreading of the 3’ss by U2AF2. In this case HNRNPA1 enables the U2AF heterodimer to reject suboptimal splice sites (Tavanez et al. 2012). In light of these diverse molecular mechanisms of HNRNPA1-dependent splicing regulation, there is a critical need to test their generality on a global scale.

In this study, we investigate how HNRNPA1 influences the association of U2AF2 with 3’ss on a transcriptome-wide scale. We used individual nucleotide resolution crosslinking immunoprecipitation (iCLIP) and high throughput sequencing to map U2AF2-RNA interactions in control or HNRNPA1 over-expression cell lines. This analysis revealed global changes in the U2AF2-RNA interactions in HNRNPA1 over-expressing cells. These experiments demonstrated that HNRNPA1 promotes loss of U2AF2 from a subset of 3’ splice sites and the concomitant increase in U2AF2 crosslinking to intronic decoy sites, including antisense Alu elements. Splicing reporter assays demonstrated that U2AF2 decoys sites are required for HNRNPA1-dependent splicing repression in HEK cells. Taken together our data suggest the intriguing hypothesis that HNRNPA1 regulates alternative splicing by altering competition between bona fide 3’ss and decoy sites for U2AF2 binding. Additionally, we discovered a role for mobile U2AF2 decoy sites in splicing regulation, suggesting that Alu elements can play a major role in the evolution of novel alternative splicing events.

## Results

To determine how HNRNPA1 influences the association of U2AF2 with 3’ splice sites on a global scale, we established an HNRNPA1-inducible expression system in HEK293 cells. We then assayed U2AF2 and HNRNPA1 protein-RNA interactions in control or HNRNPA1 over-expression cells using individual nucleotide resolution crosslinking immunoprecipitation and high throughput sequencing (iCLIP-seq) (Konig et al. 2010). We favored the over-expression approach because HNRNPA1 protein levels are elevated in many human cancers (Pino et al. 2003; Ushigome et al. 2005; Chen et al. 2010; Loh et al. 2015; Yu et al. 2015). Induction of HNRNPA1 results in an approximately 2 fold increase compared to the endogenous protein (and relative to EWSR1) and has no appreciable effect on SRSF1 or U2AF2 steady state protein levels (Fig. 1A). Additionally, the expression and localization of hnRNPC are unaffected by HNRNPA1 over-expression (Supplemental Fig. 1). We used iCLIP to purify HNRNPA1-, SRSF1-, and U2AF2-RNA complexes from control and HNRNPA1 over-expressing cells (Fig. 1B and supplemental Fig 2A and B). In all cases, the immunoprecipitated material was both UV- and antibody-dependent, nuclease sensitive and produced robust sequencing libraries (Supplemental Tables 1-3).

**Fig. 1.**
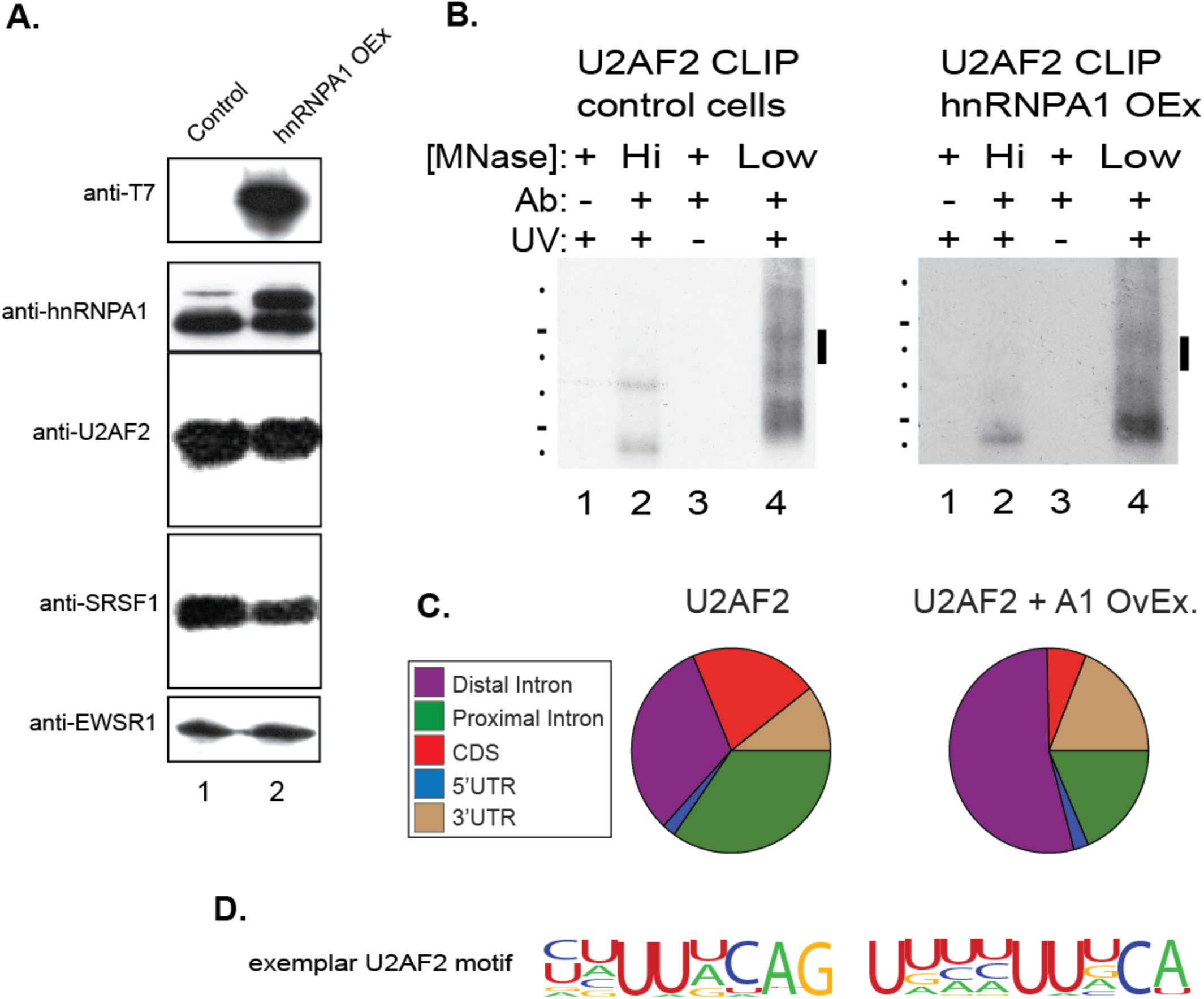
Crosslinking immunoprecipitation of U2AF2 under HNRNPA1 modulation. **(A)** Western blot analysis of SRSF1, U2AF2, HNRNPA1, the T7 epitope and EWSR1 in control (lane 1) and HNRNPA1 over-expression HEK293 cells. **(B)** Examples of iCLIP autoradiographs for U2AF2 either control or following over-expression of HNRNPA1. Protein-RNA complex shifts are UV-, antibody- and Micrococcal nuclease-sensitive. Bars denote the region of nitrocellulose blot excised for RNA isolation for iCLIP library preparation. Micrococcal Nuclease treatment at 15U (high) and 0.015U (low). **(C)** CLIPper analysis of iCLIP RNA distribution for U2AF2 in control and HNRNPA1 over-expression conditions. **(D)** Top HOMER consensus binding motifs for U2AF2 in control and HNRNPA1 over-expression conditions.

**Fig. 2.**
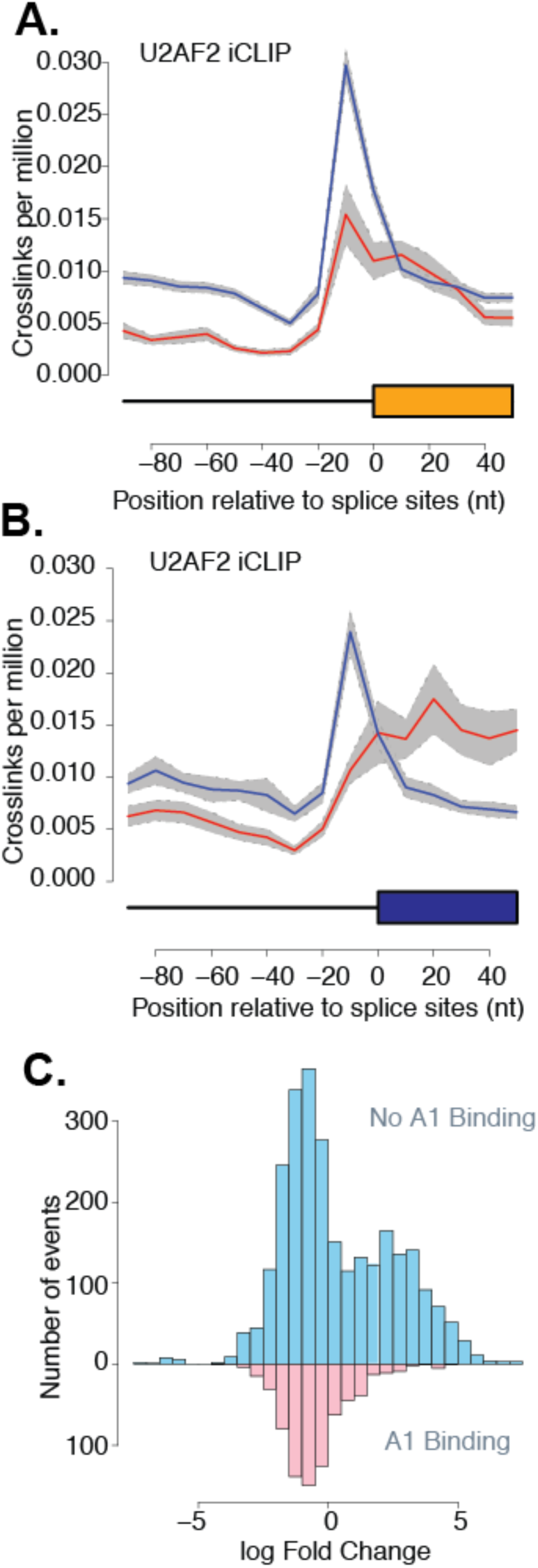
HNRNPA1 induced redistribution of U2AF2 crosslinking near 3’ splice sites. **(A, B)** Normalized crosslinking distribution for U2AF2 in wild-type (blue line) and HNRNPA1 over-expression cell lines (red line) with 95% confidence interval (gray area). Data is divided between constitutive (A) and cassette (B) exons. **(C)** Natural log fold change distribution of U2AF2 within 200bp intron regions near 3’ splice sites of cassette exons. Blue bars correspond to annotated alternative splicing events with no evidence of HNRNPA1 crosslinking in either condition and pink represents annotated events with detectable HNRNPA1 crosslinking.

After identification of binding site peaks using CLIPper (Lovci et al. 2013) (Supplemental Table 4), we observed differences in the distribution of U2AF2 peaks across genomic locations between control and HNRNPA1 over-expression cell lines (Fig. 1C). Most notably, the proportion of peaks located in coding exons (CDS) or in exon-proximal intronic regions was reduced whereas intronic peaks located more than 500 nt from exons (distal intron) increased. A similar trend was observed for SRSF1 peaks (Supplemental Fig. 2).

To determine whether HNRNPA1 over-expression influences the RNA binding specificity of U2AF2, we searched for over-represented RNA sequences within the binding site peaks (Fig. 1D). In both control cells and HNRNPA1 over-expression cells, U2AF2 peaks are characterized by a pyrimidine-rich motif, closely resembling authentic 3’ splice sites. Although the sequences are distinct from each other, SRSF1 and HNRNPA1 motifs also appear to be similar between control and HNRNPA1 over-expression cells (Supplemental Figs. 2A and B). These data suggest that over-expression of HNRNPA1 alters U2AF2 RNA-targeting with only modest effect on sequence specificity.

HNRNPA1 influences association of splicing factors such as U2AF2 with 3’ splice sites (Zhu et al. 2001; Tavanez et al. 2012). To test this hypothesis, we determined how titration of HNRNPA1 affected the distribution of SRSF1- and U2AF2-RNA cross-links relative to 3’ splice sites of constitutive or alternative cassette exons. As suggested by the peak analysis (Supplemental Fig. 3), there are no differences in HNRNPA1 crosslinking sites between control and over-expression cells (Supplemental Fig 4A and B, left panels, blue and red lines respectively). SRSF1 crosslinking to exonic sequences was modestly reduced in the HNRNPA1 over-expression cells compared to control, but the positional distribution of the SRSF1 sites relative to the 3’ss was largely unchanged for constitutive and skipped exons (Supplemental Fig 4A and B right panels, blue and red lines, respectively). U2AF2 crosslinking distribution relative to the 3’ss was substantially altered in HNRNPA1 over-expressing cells compared to the control, where a characteristic peak is observed over the 3’ss of both constitutive and skipped exons (Fig. 2A and B, blue line). By contrast, in cells overexpressing HNRNPA1, U2AF2 crosslinking density near alternative exons shifts downstream of the 3’ss and the peak is substantially reduced (Fig. 2B right panel, red line). To further determine if there is a direct relationship between HNRNPA1 binding and changes in U2AF2 or SRSF1 association with transcripts, we examined regions flanking the 3’ss of skipped exons with HNRNPA1 crosslinking in either condition (Fig. 2C; Supplemental Fig 4, Supplemental Tables 5 and 6). In regions with no detectable HNRNPA1 crosslinks, the change in U2AF2 crosslinking exhibits a bimodal distribution, which corresponds to regions flanking the 3’ss that show either increased or decreased U2AF2 crosslinking in HNRNPA1 over-expression cells relative to control cells (Fig. 2C, blue). By contrast, U2AF2 crosslinking to the vicinity of the 3’ss is significantly reduced when a direct association of HNRNPA1 is also evident (Fig. 2C, pink). For example, in both SRSF6 (serine/arginine-rich splicing factor 6 or SRp55) and PIEZO1 (Piezo-type mechanosensitive ion channel component 1) we found that U2AF2 crosslinking near the 3’ splice sites is reduced in the cell lines over-expressing HNRNPA1 (Supplemental Figure 5).

**Figure 3.**
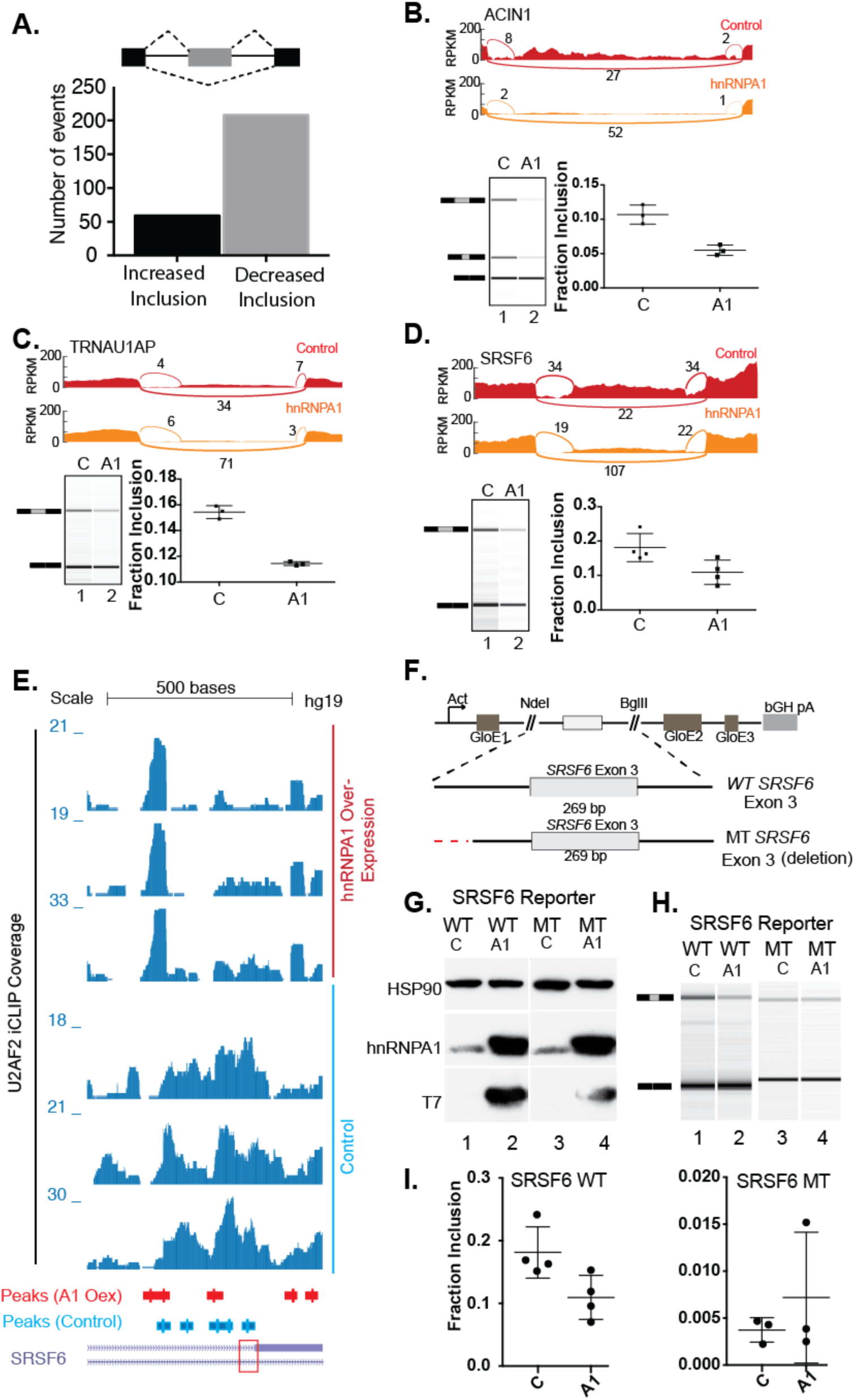
Non-canonical U2AF2 binding sites influence HNRNPA1-dependent exon skipping. **(A)** Bar graph of all HNRNPA1-dependent splicing changes as detected by MISO analysis. Events determined from the comparison of RNA-seq data derived from HNRNPA1 over-expressing HEK293T cell vs control HEK293T cells. **(B-D)** Sashimi plots showing read and junction coverage for three genes: ACIN1, SRSF6 and TRNAU1AP. Virtual Gel representation of Agilent 2100 Bioanalyzer DNA 1000 assay of RT-PCR products corresponding to endogenous ACIN1, SRSF6 and TRNAU1AP from control and HNRNPA1 over-expression cells (lower left panel). Bar graph depicting mean exon inclusion for TRNAU1AP and SRSF6 splicing reporters quantified using an Agilent 2100 Bioanalyzer with standard deviation bars. *P*< 0.05, *; *P*< 0.01, **; *P*< 0.001, *** (lower right panel). **(E)** UCSC Genome Browser screen shot of U2AF2 iCLIP read coverage at the SRSF6 exon 3 locus in control or HNRNPA1 overexpression cells (bottom and top three tracks, respectively). Peaks called by CLIPPER are depicted below the coverage tracks. Red box denotes the 3’ss of SRSF6 exon 3. **(F)** Splicing reporter constructs created representing wild-type (Wt) or mutant (Mt) versions of alternative exons in SRSF6. Non-canonical U2AF2 binding sites in SRSF6 were mutated (red lines) by deletion. GloE1, GloE2, and GloE3 designate exons 1–3 of beta-globin. The polyadenylation signal from the bovine growth hormone 1 gene is indicated by bGH pA. **(G)** Representative western blots probed with antibodies against HSP90, HNRNPA1 and the T7 epitope. Lysates were prepared from HEK293T cells transiently transfected with either wild type (WT) or mutant (MT) SRSF6 reporter construct and control or T7 epitope tagged HNRNPA1. **(H)** Virtual Gel representation of Agilent 2100 Bioanalyzer DNA 1000 assay of RT-PCR products from SRSF6 splicing construct transfections. **(I)** Bar graph depicting mean exon inclusion for SRSF6 splicing reporters quantified using an Agilent 2100 Bioanalyzer with standard deviation bars. *P*< 0.05, *.

**Fig. 4.**
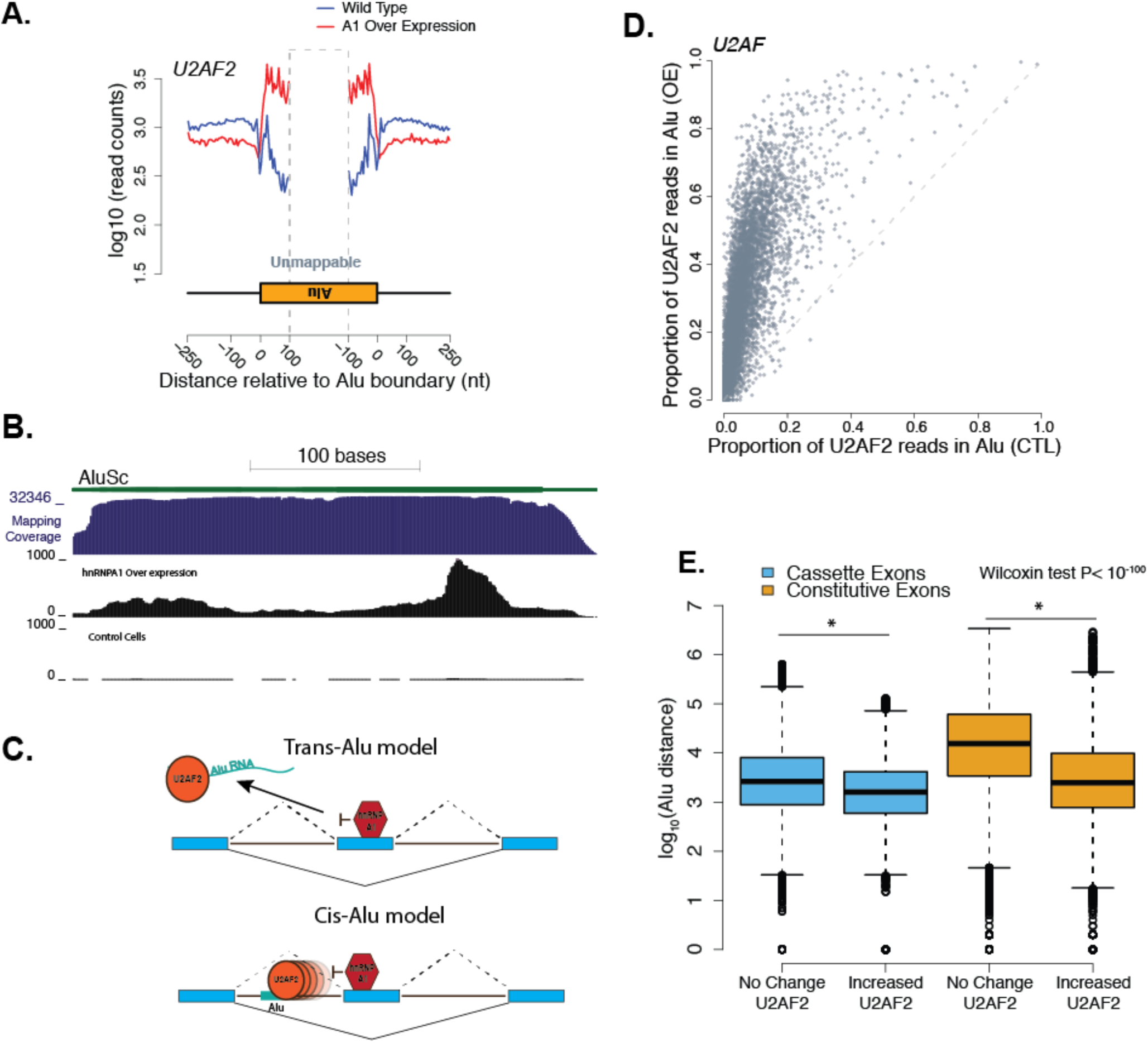
HNRNPA1 over-expression correlates with global redistribution of U2AF2 signal to *Alu* RNA elements. **(A)** Aggregated crosslinking sites on *Alu* elements and nearby regions for U2AF2. Blue represents wild-type binding of the given RNA binding protein and red represents HNRNPA1 over-expression of the log_10_ number of iCLIP read counts across all antisense-Alu elements. **(B)** Distribution of aggregated U2AF2 iCLIP peaks on *Alu* subtype *AluSc* under control and HNRNPA1 over-expression conditions. **(C)** Model representing two potential modes by which U2AF2 may associate with Alu RNA: *trans*-competition suggests U2AF2 binds to Alu elements on other RNAs, while a *cis*-competition suggest U2AF2 binds to *Alu* elements within the same RNAs that a particular exon is associated. **(D)** Scatter plot of all human cassette exons measuring the proportion of U2AF2 iCLIP crosslinking sites found within *Alu* elements within the cassette exon event over the total number of crosslinks found within the event. Proportions from control and HNRNPA1 over-expression samples are compared for each individual cassette exon event. **(E)** Box plot representing the distance of *Alu* elements from cassette exons (blue) and constitutive exons (orange) that show no change in U2AF2 cross-linking versus those that show an increase in U2AF2 crosslinking.

To determine if changes in U2AF2 crosslinking correlated with HNRNPA1-dependent splicing regulation, we sequenced poly-A+ selected RNA libraries from control and HNRNPA1 over-expression cells (supplemental table 7). Of the 267 HNRNPA1-regulated cassette exons, the majority exhibited increased levels of exon skipping upon HNRNPA1 over-expression (Fig. 3A, Supplemental Table 8). 83 of the 267 differentially spliced exons also had detectable U2AF2 crosslinking in either cell line. Of those 83 exons, ∼90% exhibited an HNRNPA1-dependent increase in exon skipping compared to ∼70% of exons with no detected U2AF2 crosslinking (Table 1, *P* < 0.0025, Fishers Exact Test). ∼54% (44/83) of HNRNPA1-regulated splicing events also exhibited changes (<2 fold) in U2AF2 intronic crosslinking (Table 2 and Supplemental Table 5 and 6), whereas 46% show HNRNPA1-dependent increases in U2AF2 crosslinking in the same region. Many of these HNRNPA1-sensitive exons exhibited redistribution of U2AF2 signal from near 3’ss to distal, upstream crosslinking sites, suggesting their possible function in HNRNPA1-mediated splicing regulation (Figure 3E and Supplemental Figs. 6-16). We use transient transfection of T7 epitope tagged HNRNPA1 and RT-PCR to validate several endogenous splicing events exhibiting HNRNPA1-dependent reduction in U2AF2-3’ss crosslinking (Fig. 3B-D, upper panels). As expected, we observed a significant reduction in exon inclusion for ACIN1, SRSF6 and TRNAU1AP in HNRNPA1 over-expression cells relative to control (Fig. 3B-D, lower panels).

**Table 1.** Summary of HNRNPA1-dependent changes in exon skipping with detectable U2AF2 crosslinking.

|  |  |  |  |
| --- | --- | --- | --- |
|  | Total # of Events | U2AF2 Crosslinking | No U2AF2 Crosslinking |
| HNRNPA1-dependent exon inclusion | 59 | 9 | 50 |
| HNRNPA1-dependent exon skipping | 208 | 74 | 134 |

**Table 2.** Summary of HNRNPA1-dependent changes in exon skipping and U2AF2 positioning.

|  |  |  |  |  |
| --- | --- | --- | --- | --- |
|  |  | HNRNPA1-dependent change in U2AF2 crosslinking near 3'ss |  |  |
|  | Total # of Events | Increased U2AF2 | Decreased U2AF2 | Redistribution to Alu |
| HNRNPA1-dependent exon inclusion | 3 | 2 | 1 | 2 |
| HNRNPA1-dependent exon skipping | 41 | 19 | 22 | 17 |

To determine if non-canonical U2AF2 binding sites are involved in HNRNPA1-dependent exon skipping we created matched pairs of beta-hemoglobin-based (*HBB1*) splicing reporter gene constructs (Rothrock et al. 2003) containing the wild-type or mutated intronic elements upstream of the HNRNPA1-sensitive exons found in the *SRSF6* pre-mRNA (Fig. 3E). The *SRSF6* splicing reporter constructs were co-transfected into HEK293T cells with epitope-tagged HNRNPA1 expression plasmid or a control plasmid (Fig. 3F). Because inclusion of the test exons may induce nonsense-mediated decay (NMD) by inducing an in-frame premature termination codon (PTC), as is the case with SRSF6 we assayed splicing in the presence of the translation inhibitor emetine dihydrochloride, a potent inhibitor of NMD *in vivo* (Noensie and Dietz 2001; Lareau et al. 2007; Ni et al. 2007). As shown in Fig. 3G, RT-PCR of wild-type SRSF6 splicing reporters show expected increases in exon skipping in response to HNRNPA1 over-expression compared to endogenous levels. Deletion of the distal intronic U2AF2 binding site attenuated HNRNPA1-dependent exon skipping. Quantification of amplicon ratios demonstrated that mutation of the putative HNRNPA1-induced U2AF2-associated intronic elements disrupt splicing sensitivity to HNRNPA1 over-expression (Fig. 3H; *P* < 0.0228 respectively, ratio paired T-test).

Previous work by Zarnack et al. demonstrated that hnRNP proteins, such as hnRNP C, can antagonize binding of U2AF2 to antisense *Alu* elements, to repress their exonization (Zarnack et al. 2013). We asked if HNRNPA1 similarly repressed U2AF2 crosslinking to antisense Alu elements by measuring the global distribution of crosslinks for each protein overlapping of antisense Alu elements throughout intronic regions with titration of HNRNPA1 levels. Surprisingly upon over-expression of HNRNPA1, we detected a dramatic increase in U2AF2 crosslinking to antisense Alu-containing RNA transcripts compared to control cells (Fig 4A). To visualize HNRNPA1-dependent accumulation of U2AF2 in Alu elements we calculated the coverage of all peaks across a consensus sequence assembled from all the Alu subtypes in the human genome (see methods). For example, for subtype *AluSc*, HNRNPA1 over-expression induces U2AF2 crosslinking near the 3’ end of the element compared to control cells (Fig. 4B). Conversely, HNRNPA1 crosslinking globally decreases over Alu elements with over-expression (Supplementary Fig 17A,C). By contrast to U2AF2, crosslinking of SRSF1 to antisense Alu elements shows no appreciable changes (Supplementary Fig. 17 B,C), suggesting that the effect of HNRNPA1 is specific to U2AF2.

To determine if Alu elements exhibiting HNRNPA1-dependent increases in U2AF2 crosslinking are located in *cis*-relative to annotated skipped exons (Alu elements upstream or downstream of a splicing event) or in *trans-* (another locus, Fig. 4C), we compared the proportion of U2AF2 crosslinks within Alu-elements relative to flanking sequences across individual exon skipping events in control or HNRNPA1 over-expression cells. The scatter plot shown in Fig 4D (data in Supplemental Tables 9 and 10), demonstrates that the proportion of U2AF2 crosslinks present in Alu elements increases significantly across virtually all exon skipping events, upon HNRNPA1 over-expression, whereas the proportion of HNRNPA1 crosslinks are decreased (Supplemental Fig. 17D; data in Supplemental Tables 11 and 12). By contrast, the proportion of SRSF1 crosslinks to Alu elements is refractory to changes in HNRNPA1 expression levels (Supplemental Fig. 17E; data in Supplemental Tables 13 and 14). These data demonstrate a global change in U2AF2-Alu association and refute the hypothesis that a few spurious Alu-elements are responsible for the signal observed in Fig 4A. To determine if *cis-*Alu elements are involved in HNRNPA1-regulated alternative splicing we manually curated the list of 83 splicing events that also had detectable U2AF2 crosslinking. We found that 41% of HNRNPA1-dependent exon skipping events exhibited redistribution of U2AF2 to adjacent Alu elements (Table 1). Taken together, these data demonstrate that over-expression of HNRNPA1 influences the association of U2AF2 with antisense Alu elements, which may contribute to splicing regulation.

Alu elements influence alternative splicing, although the mechanisms are poorly understood (Sorek et al. 2002; Lev-Maor et al. 2008; Gal-Mark et al. 2009; Schwartz et al. 2009; Pastor and Pagani 2011). Splicing regulatory elements often exhibit position-dependent and context-dependent functions (Fu and Ares 2014). To determine if Alu-elements with HNRNPA1-dependent changes in U2AF2 crosslinking exhibit positional bias we measured their distance relative to the 3’ splice site of constitutive or skipped exons. In general, we observed that Alu elements are closer to skipped than constitutive exons (*P* < 1.4e-47, Wilcoxon rank-sum test, Fig. 4E, compare blue and orange boxes). But yet, those Alu elements with HNRNPA1-dependent increases in U2AF crosslinking are significantly closer to exons than those that are unchanged (*P* < 9.5e-93, Fig 4E). Taken together our data suggest the intriguing hypothesis that Alu-elements may function as *cis*-regulatory elements that compete with authentic exons for binding to splicing factors.

## Discussion

HNRNPA1 represses splicing through diverse mechanisms. Our data suggest that HNRNPA1 alters the competition between bona fide and decoy 3’ splice sites. Because HNRNPA1 forms a ternary complex with U2AF and the 3’ss, HNRNPA1-dependent redistribution of U2AF2 from bona fide splice sites to “decoy” sites might arise from direct competition for these splicing factors at 3’ splice sites. In cells overexpressing HNRNPA1 we observed a pronounced shift in U2AF2 binding sites relative to control cells. Based on U2AF2 peak distributions, this HNRNPA1-dependent redistribution involves loss of proximal-intron and recognition of distal-intronic peaks (Fig. 1). A similar pattern was observed at the single nucleotide level where U2AF2 crosslinking density near alternative 3’ splice sites was reduced in HNRNPA1 over-expression cells and distributed across exons (Fig. 2). Despite evidence for global changes in U2AF2 binding position, we found nearly identical sequence motifs enriched at peaks in both cell lines suggesting little, if any, differences in U2AF2 RNA binding specificity (Fig. 1). Splicing reporter assays revealed that upstream polypyrimidine tracks identified by iCLIP were necessary for HNRNPA1-dependent splicing regulation (Fig. 3). These data also support the hypothesis that distant U2AF2 binding sites are likely to function as splice site decoys that contribute to HNRNPA1-dependent splicing silencing.

Our findings are well-aligned with a recent census of U2AF2 binding sites in HeLa cells, which documented a position-dependent code for U2AF2 in splicing regulation (Shao et al. 2014). Perhaps most relevant to the work presented here is their observation that non-canonical U2AF2 binding sites located upstream or within alternative exons correlates with exon skipping (Shao et al. 2014; Cho et al. 2015). Our results suggest a role for HNRNPA1 in promoting recognition of non-canonical sites by U2AF2. Our data (Supplemental Fig 5), as well as protein-protein interaction screens (Akerman et al. 2015), argue against a direct interaction between HNRNPA1 and U2AF2. One potential model to explain our results is that HNRNPA1 and the U2AF heterodimer compete for binding to the 3’splice site, resulting in displacement of U2AF (Zhu et al. 2001; Okunola and Krainer 2009; Jain et al. 2017). Alternatively, the interaction of HNRNPA1 with nearby positions may alter RNA conformation thereby interfering with U2AF-3’ss association.

The results presented here may also illuminate the role of HNRNPA1 and U2AF in proofreading of the 3’ss (Tavanez et al. 2012). Tavarnez et al demonstrated the HNRNPA1 is required for discrimination of AG and CG dinucleotides at 3’splice sites. HNRNPA1 enables U2AF to ignore the non-canonical CG-containing splice sites but to productively engage AG-containing splice sites. We observed a global decrease in U2AF2 crosslinking near alternative 3’ss, but little change near constitutive splice sites. Given that alternative exons are typically flanked by weak splice sites, it is possible that this proof reading mechanism also contributes to regulation of alternative splicing. Tavarnez et al. also noted that depletion of HNRNPA1 from HeLa cells resulted in increased U2AF association at spurious sites, suggesting a lack of fidelity. Their work identified several sites within the 3’UTRs of intronless messages that accumulate U2AF when HNRNPA1 is knocked down. We also observed a global increase in U2AF2 crosslinking to 3’UTRs but not CDS regions when HNRNPA1 is over expressed (supplemental table 3). It will be important to understand the RNA determinants (sequence, structure, splice site strength) that distinguish between a proofreading function for HNRNPA1 and a simple competition with U2AF (Jain et al. 2017).

Alu elements influence gene expression in diverse ways (Hasler and Strub 2006; Chen and Carmichael 2009; Gong and Maquat 2011; Pastor and Pagani 2011; Kelley et al. 2014; Tajnik et al. 2015). Recently, Zarnack et al. demonstrated that hnRNP C competes with U2AF2 to repress inclusion of Alu-derived exons in mRNA (Zarnack et al. 2013). We find that HNRNPA1 over-expression correlates with increased U2AF2 association with Alu-derived RNA sequences. We did not observe any change in hnRNP C protein expression or localization (Supplemental Fig. 1). Taken together, our data demonstrate that Alu-derived sequences function as RNA regulatory elements that respond to changes to the intracellular concentration of splicing factors. It is intriguing to speculate that, as primate-specific elements, antisense Alu elements may influence the evolution of splicing regulation by modulating recognition of bona fide exons. Our results suggest the intriguing hypothesis that antisense Alu elements contribute to species-specific differences in alternative splicing throughout the primate lineage.

## Materials and Methods

### iCLIP analysis of U2AF2, SRSF1 and HNRNPA1

iCLIP was performed as previously described (Konig et al. 2010; Huppertz et al. 2014). Briefly, TREX FLP-in HEK293T cells (Invitrogen) lacking or containing a stable, inducible T7-tagged version of HNRNPA1. Cells were treated with tetracycline for 24 hr and then irradiated with UV-C light to form irreversible covalent cross-link between proteins and nucleic acids *in vivo*. After cell lysis, RNA was partially fragmented using low concentrations of Micrococcal nuclease, and U2AF65-, SRSF1-, or HNRNPA1–RNA complexes were immunopurified with U2AF65, (MC3;SCBT), SRSF1 (96;SCBT), and HNRNPA1 (4B10;SCBT) antibodies immobilized on protein A–coated magnetic beads (Life Technologies), respectively. After stringent washing and dephosphorylation (Fast AP, Fermentas), RNAs were ligated at their 3′ ends with a pre-adenylated RNA adaptor (Bioo Scientific) and radioactively labeled to allow visualization. Samples were run using MOPS-based protein gel electrophoresis (in-house recipe) and transferred to a nitrocellulose membrane. Protein-RNA complexes migrating 15-80 kDa above free protein were cut from the membrane, and RNA was recovered from the membrane by proteinase K digestion under denaturing (3.5 M Urea) conditions. The oligonucleotides for reverse transcription contained two inversely oriented adaptor regions adapted from the Bioo NEXTflex small RNA library preparation kit (Bioo Scientific), separated by a BamHI restriction site as well as a barcode region at their 5′ end containing a 4-nt experiment-specific barcode within a 5-nt random barcode to mark individual cDNA molecules. cDNA molecules were size-purified using denaturing PAGE gel electrophoresis, circularized by CircLigase II (Epicenter), annealed to an oligonucleotide complementary to the restriction site and cut using BamHI (NEB). Linearized cDNAs were then PCR-amplified using (Immomix PCR Master Mix, Bioline) with primers (Bioo) complementary to the adaptor regions and were subjected to high-throughput sequencing using Illumina HiSeq. A more detailed description of the iCLIP protocol has been published (Huppertz et al. 2014)

### Mapping and analysis of iCLIP sequencing data

Single-end reads generated by Illumina HiSeq were inspected for the presence of adaptor sequences. Reads containing sequences corresponding to the 3’RNA adaptor were retained if they were at least 30bp long after the adaptor sequence was trimmed off. The first 9bp in each read from the iCLIP library preparation, containing an internal barcode comprising 4bp for replicate identification and 5bp of random nucleotides for use in duplicate mapping removal, were also removed before mapping. Trimmed reads were checked for mapping to a repeat filter comprising RepeatMasker elements in the human genome using Bowtie2 (Langmead et al. 2009). Reads that passed the repeat filter were mapped to the transcriptome and genome with Tophat (Kim et al. 2013). If reads mapped equally well to multiple loci, a single mapping was selected randomly by Tophat. Duplicate mappings from each replicate were reduced to one per position if they had the same genomic endpoints and if they originated from reads with the same set of random 5bp nucleotides. Following mapping and duplicate removal, individual reads were truncated to their 5’ ends to represent the site of crosslinking consistent with the iCLIP methodology. For all samples only such crosslinking sites found to have non-zero mapping counts in two out of three replicates (or two out of two duplicates where applicable) were considered to be biologically reproducible candidates for further analysis. The counts at such reproducible crosslinking sites were summed over all replicates to create an aggregated dataset for each cell condition and CLIP. To determine background from the iCLIP datasets, the two cell conditions (control and HNRNPA1-overexpressing) were temporarily further aggregated for each CLIP (U2AF, SF2, A1) and those binding sites that had non-zero counts in all three temporary aggregate datasets were determined. A 41 nt mask was created by extending 20nt upstream and 20nt downstream from each such 3-way common binding site. The aggregated data set of binding sites for each cell condition and CLIP was then filtered using this mask, keeping only sites outside the mask that also had a mapping count of at least 3 in the aggregate data. These aggregated and filtered data were used for downstream analyses. This aggregation and filtering strategy was adapted from previously described iCLIP analysis pipelines (Friedersdorf and Keene 2014; Flynn et al. 2015). For use as input to CLIPper (Lovci et al. 2013), the filtered (single nucleotide) binding sites were expanded by 15nt upstream and 15nt downstream.

CLIPper (CLIP-seq peak enrichment; https://github.com/YeoLab/CLIPper) was used to determine genomic distribution of RNA crosslinking peaks as well as identify clusters representing binding sites for HNRNPA1, U2AF2, and SRSF1 for each condition as previously described (Lovci et al. 2013). For each condition, the resulting iCLIP peak data of the replicates were merged. The peaks were annotated to the human genome (hg19) and then divided into categories based on their genomic locations including CDS, intron, and UTR. The peaks in each category were further subsetted based on whether they overlapped with Alu elements. For the motif analysis, 50bp sequences were extracted from the peak regions (crosslinking site ± 25bp). Strand-specific MEME-ChIP and HOMER analyses were performed on these sequences to find 6-10bp long, enriched motifs.

### RBP Binding Analysis

40,769 cassette exons were extracted from MISO human genome (hg19) alternative events annotation version 2. 200,880 constitutive exons were extracted from RefSeq gene annotation by excluding the exons that overlap with cassette exons. Gene differential expression analysis was performed using edgeR. 40,952 constitutive exons that were not significantly differentially expressed (FDR > 0.05) were used in further analysis.

For each RNA-binding protein in each cell line, the iCLIP reads of all the replicates were merged together (Flynn et al. 2015). The start positions of the reads were considered as crosslinking sites. The number of reads near the 3’ splice site (100bp into the intron, 50bp into the exon) of each exon were calculated based on a 10bp window. The raw read counts were normalized by the total library size.

The changes in binding of U2AF2 and SF2 near 3’ splice sites were further analyzed with edgeR. Read counts were calculated for 200bp intron regions near the 3’ splice sites of the cassette exons. For each RBP, the regions with more than one count per million (CPM) in at least half of the replicates in either of the cell line were used for binding change analysis.

### RBP Binding Near Alu Elements

315,974 antisense Alu elements were extracted from RepeatMasker. The merged iCLIP data for each condition was down-sampled to 1M reads. The total number of sense strand reads were calculated for Alu and nearby regions (250bp from Alu boundary). For each cassette exon events (cassette exon + up/downstream introns + up/downstream exons), the number of reads in antisense Alu elements, and the total number of reads in the whole event were calculated separately. The proportion of reads that fall into antisense Alu elements for each event was used to represent the RBP binding change in Alu regions.

### mRNA-seq of control or HNRNPA1-overexpressing HEK293 cells

RNA was isolated from whole cell lysates of control and HNRNPA1-overexpressing TREX Flp-IN HEK 293T cells using TRI-Reagent LS (Sigma). Poly-A+ sequencing libraries were prepared using the TrueSeq RNA library prep kit (Illumina, San Diego, CA). Each condition was analyzed in duplicate using the HiSeq2000.

### Quantification of alternative splicing by RNA-Seq

Poly-A+ transcriptome sequencing reads were mapped to the human reference genome (hg19) with TopHat2. Mapped reads of duplicates were merged together for splicing analysis. Splicing change was analyzed with MISO (Katz et al. 2010). The MISO result was filtered with the following parameters: --num-inc 1 --num-exc 1 --num-sum-exc 10 --delta-psi 0.20 --bayes-factor 10. After filtering, 267 skipped exon events were left for further analysis.

### Mapping to an ALU consensus sequence

ALU element annotations were obtained from the hg19 UCSC Genome Browser RepeatMasker track. A strategy similar to the one described in Jacobs et al. (Jacobs et al. 2014) was used to show the CLIPper peak density over all ALU elements as follows: after removal of the longest 2%, the top 50 longest human ALU sequences were aligned with MUSCLE and used to construct a consensus sequence. CLIPper peaks were mapped from the genomic position to the consensus sequence position using a BLAT alignment of the repeat to the consensus and the coverage of summits per bp of the consensus AluSc was plotted in Supplemental Figure 5c. A Genome Browser Session displaying the Repeat Masker Data can be found here: (http://genome.ucsc.edu/cgi-bin/hgTracks?hgS_doOtherUser=submit&hgS_otherUserName=Max&hgS_otherUserSessionName=pubRepeats2Sa_nford).

## Abbreviations

3’ss: 3’ splice site
iCLIP: individual nucleotide resolution cross-linking immunoprecipitation
HNRNPA1: heterogeneous nuclear ribonucleoproteins A
U2AF2: U2 Small Nuclear RNA Auxiliary Factor 2
SRSF1: Serine/arginine-rich splicing factor 1
SRSF6: Serine/arginine-rich splicing factor 6
PIEZO1: Piezo type mechanosensitive ion channel component 1
TRNAU1AP: tRNA selenocysteine 1-associated protein 1
hnRNP C: heterogeneous nuclear ribonucleoproteins C1/C2

## Declarations

## Ethics approval and consent to participate

Ethics approval was not needed for this study.

## Consent for publication

Not applicable.

## Availability of data and materials

The datasets generated and analysed during the current study are available on GEO (Gene Expression Omnibus; https://www.ncbi.nlm.nih.gov/geo/) under accession GSE83923.

## Authors contributions

JMH, HL, YL and JRS designed the experiments. JMH, HL, GK, JMD, and SK performed the experiments. JMH, HL, GK, JMD, YL and JRS wrote and edited the manuscript. All authors read and approved the final manuscript.

## Acknowledgements

We thank Professors Javier Caceres (MRC HGU, Edinburgh), Manny Ares (UCSC) and Al Zahler (UCSC) for discussion of the project. We thank Julia Philipp (UCSC) for thoughtful comments on the manuscript.

## Supplementary Figure Legends

**Fig S1.**
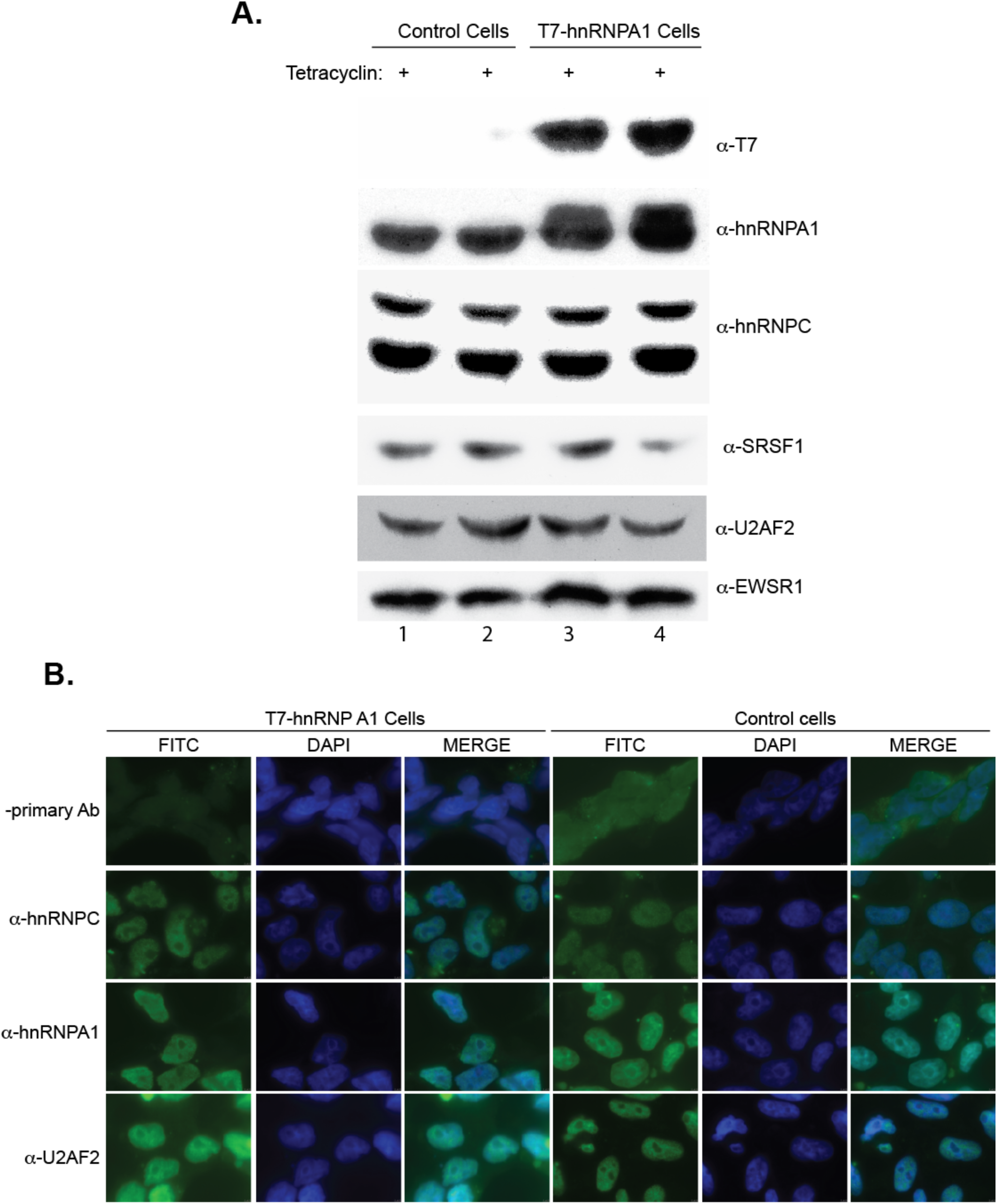
Overexpression of hnRNP A1 does not confer appreciable changes in hnRNP C1/C2 expression in HEK293T cells. **(A)** Western blot of TREX HEK293T control cells and those containing a tetracycline-inducible T7-tagged version of hnRNP A1 (T7-A1). After 24 h cell lysate was subjected to immunoblotting for hnRNP A1, T7 protein tag, hnRNP C1/C2, and EWS as a loading control. Experiment was performed in duplicate. **(B)** Intracellular distribution of hnRNP C, hnRNP A1, and U2AF2 analyzed by immunofluorescence. Control and T7-hnRNP A1 HEK293T TREX Flp-In were treated with tetracycline, fixed with 4% paraformaldehyde. Fixed cells were stained with anti-hnRNP C (4F4, Santa Cruz Biotechnology), anti-hnRNP A1 (4B10, Santa Cruz Biotechnology), or anti-U2AF2 (MC3, Santa Cruz Biotechnology) antibodies. Cells were subsequently stained with Cy2-conjugated goat anti-mouse antibody and co-stained with DAPI for nuclear reference.

**Fig. S2.**
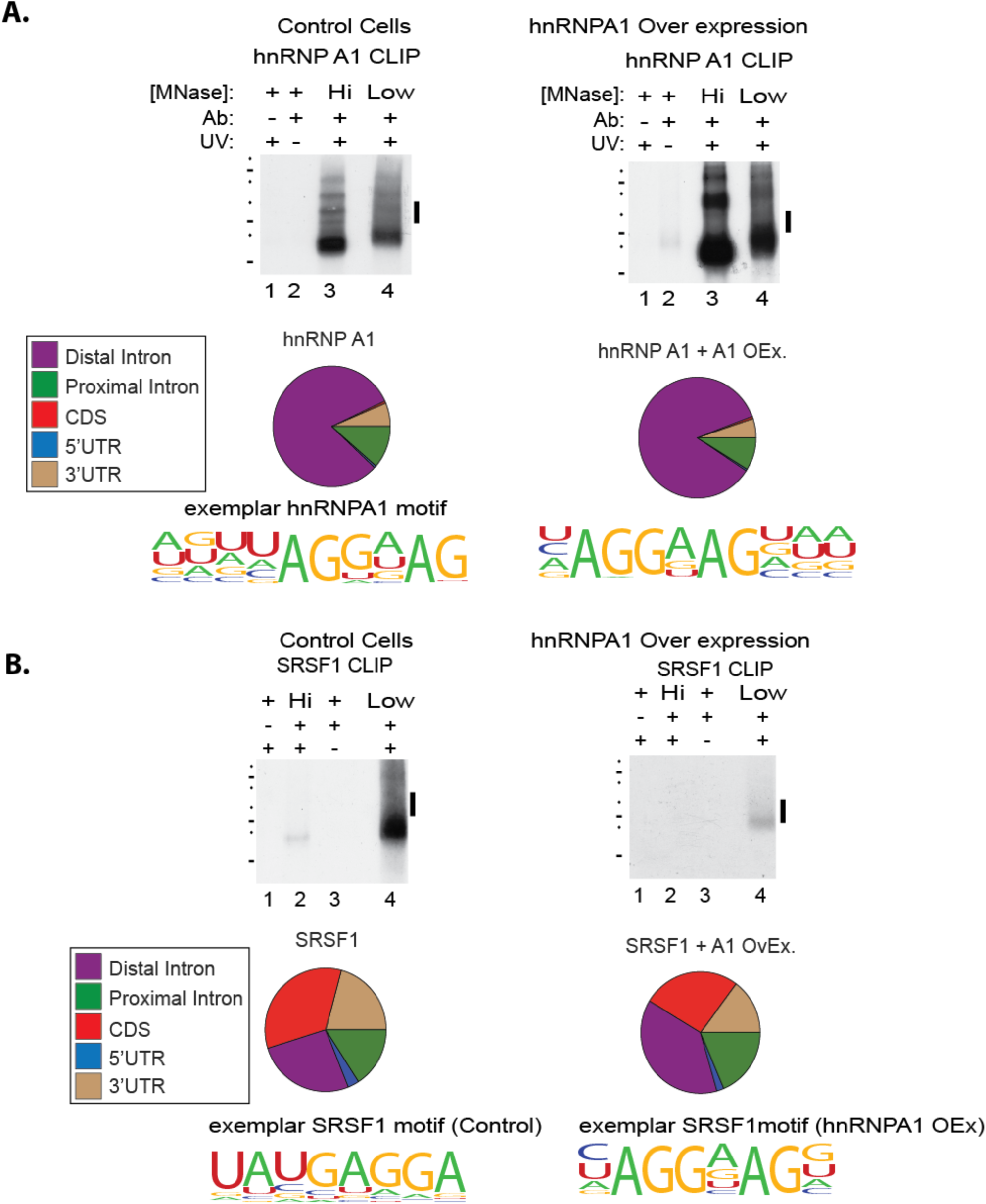
Crosslinking immunoprecipitation of hnRNPA1 and SRSF1 under hnRNPA1 modulation. **(A)** Summary of iCLIP of hnRNPA1 under control and hnRNPA1 overexpression conditions. Examples of iCLIP autoradiographs for hnRNPA1 under control and overexpression of hnRNPA1. Protein-RNA complex shifts are UV-, antibody- and Micrococcal nuclease-sensitive. Bars denote the region of nitrocellulose blot excised for RNA isolation for iCLIP library preparation. CLIPper analysis of iCLIP RNA distribution for U2AF2 in control and hnRNPA1 overexpression conditions. Top HOMER consensus binding motifs for hnRNPA1 in control and hnRNPA1 overexpression conditions. **(B)** Top left panel, autoradiograph from SRSF1 IP from control cells. Top left panel, autoradiograph from SRSF1 IP from hnRNP A1 over expression cells. Middle left panel, annotation of SRSF1 peaks identified by CLIPPER and consensus motifs identified by HOMER from control cells. Middle right panel, annotation of SRSF1 peaks identified by CLIPPER and consensus motifs identified by HOMER from hnRNP A1 over expression cells.

**Fig. S3.**
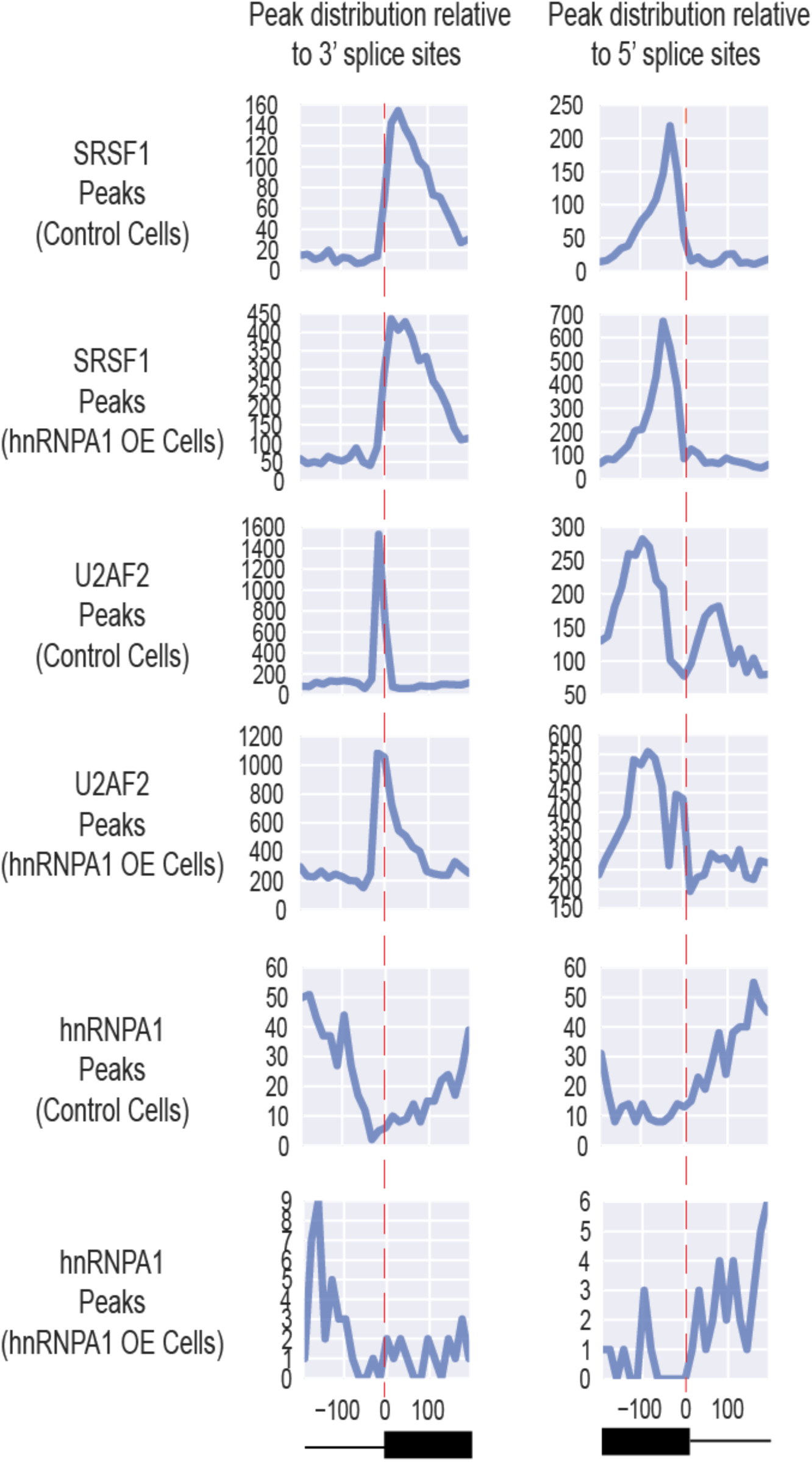
Distribution of hnRNPA1, SRSF1 and U2AF2 peaks relative to splice sites. The frequency of peaks occurring at different positions relative to splice sites is shown.

**Fig S4.**
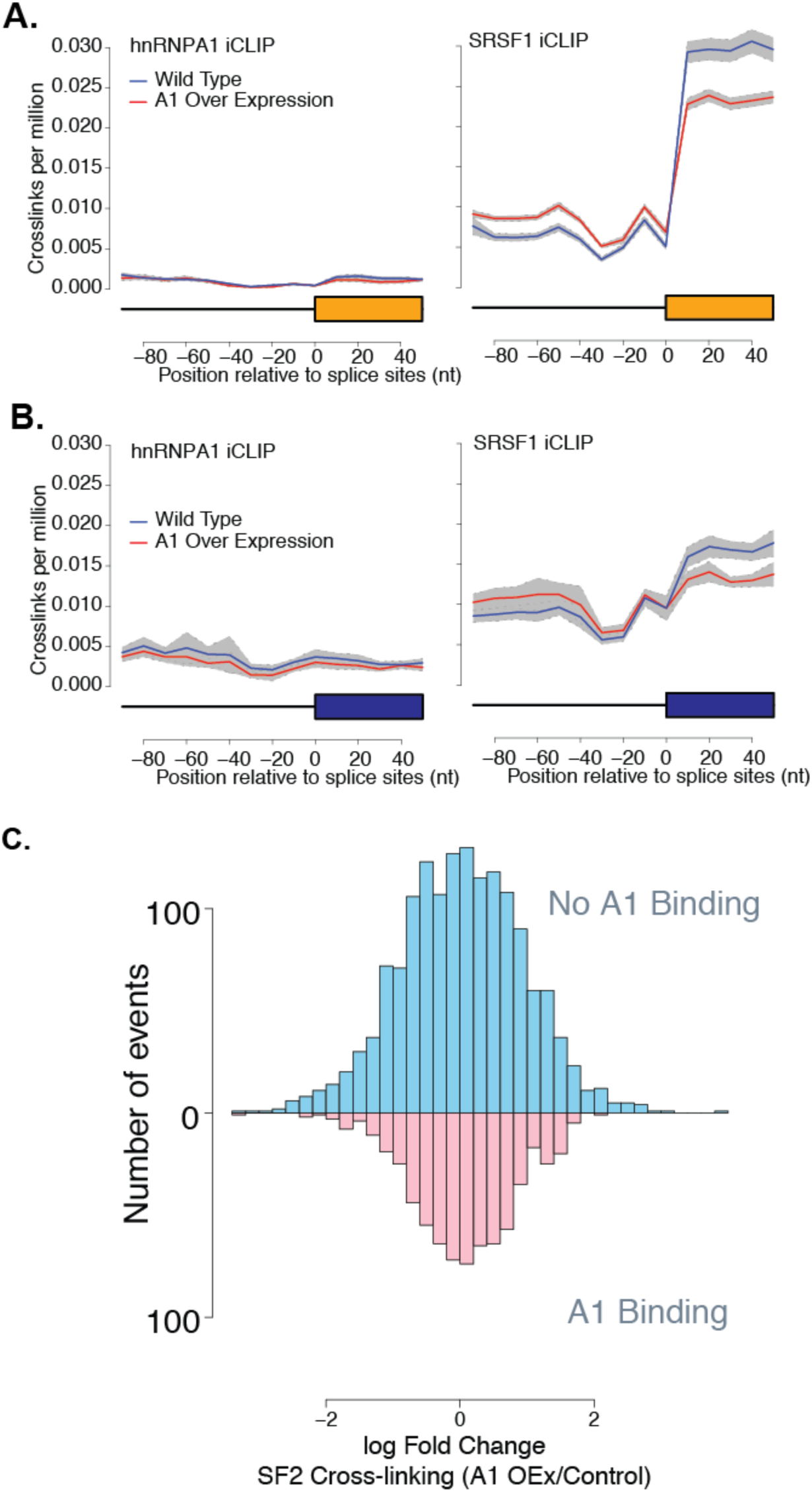
Global hnRNP A1 and SRSF1 crosslinking remains constant near 3’ splice sites under hnRNPA1 overexpression. **(A,B)** Normalized crosslinking distribution for hnRNPA1 (left panel), SRSF1 (right panel) in wild type (blue line) and hnRNPA1 overexpression cell lines (red line) with 95% confidence interval (grey area). Data is divided between constitutive (A) and cassette (B) exons. **(C)** Natural log fold change distribution of SRSF1 within 200bp intron regions near 3’ splice sites of cassette exons. Blue bars correspond to annotated alternative splicing events with no evidence of hnRNPA1 crosslinking in either condition and pink represents annotated events with detectable hnRNP A1 crosslinking.

**Fig. S5.**
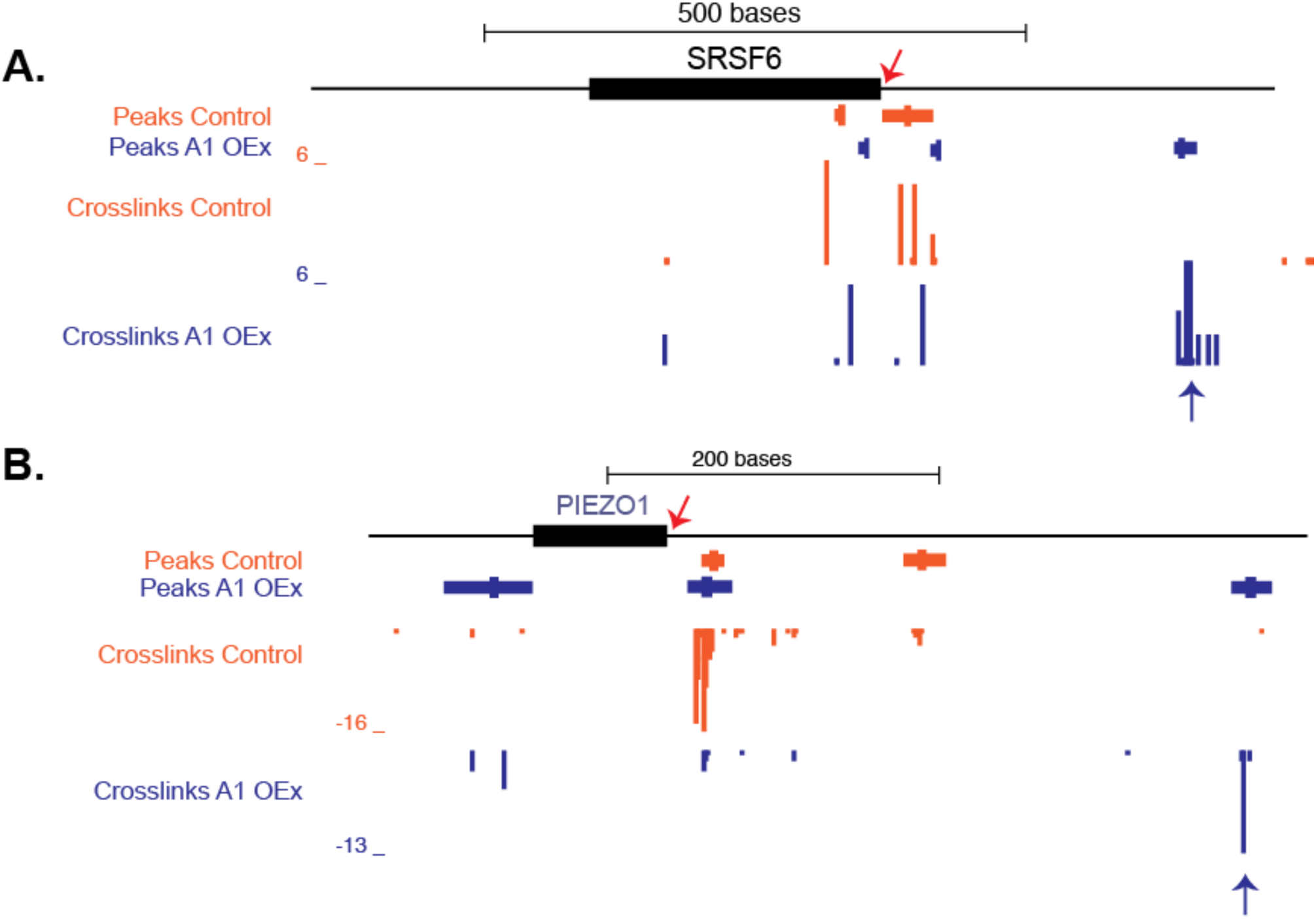
Examples of hnRNPA1-dependent modulation of U2AF2 crosslinking. UCSC genome browser examples of two genes SRSF6 (A) and PIEZO1 (B) and iCLIP crosslinking site (5’ends of reads) coverage data for U2AF2 under control and hnRNPA1 overexpression. Alternative splicing changes can be observed in figure 3 (SRSF6) and supplemental figures 10 and 13 (PEIZO1 and SRSF6, respectively).

**Fig S6.**
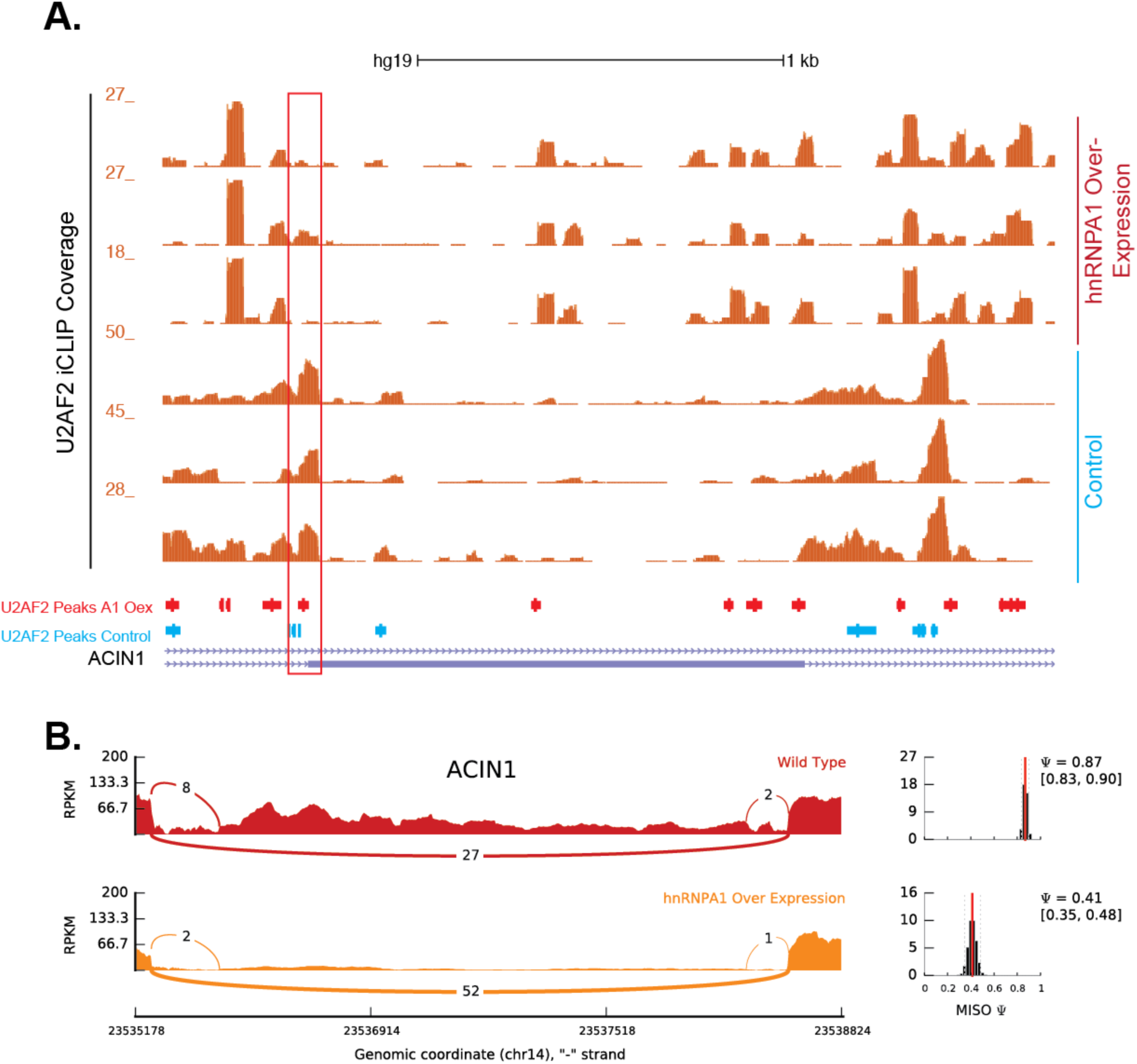
hnRNPA1-dependent modulation of U2AF2 crosslinking and alternative splicing of ACIN1 locus. **(A)** UCSC genome browser of ACIN1 alternative exon and CLIP read coverage data for U2AF2 under control (blue text) and hnRNPA1 overexpression (red text). Red box highlights region in which U2AF2 is seen to redistribute between two conditions. **(B)** Sashimi plots showing read and junction coverage for ACIN1 gene under wildtype (red) and hnRNPA1 over-expression (orange) conditions. PSI (percent splicing index) values are provided for each condition reflecting inclusion level of exon shown in genome browser (A) image.

**Fig S7.**
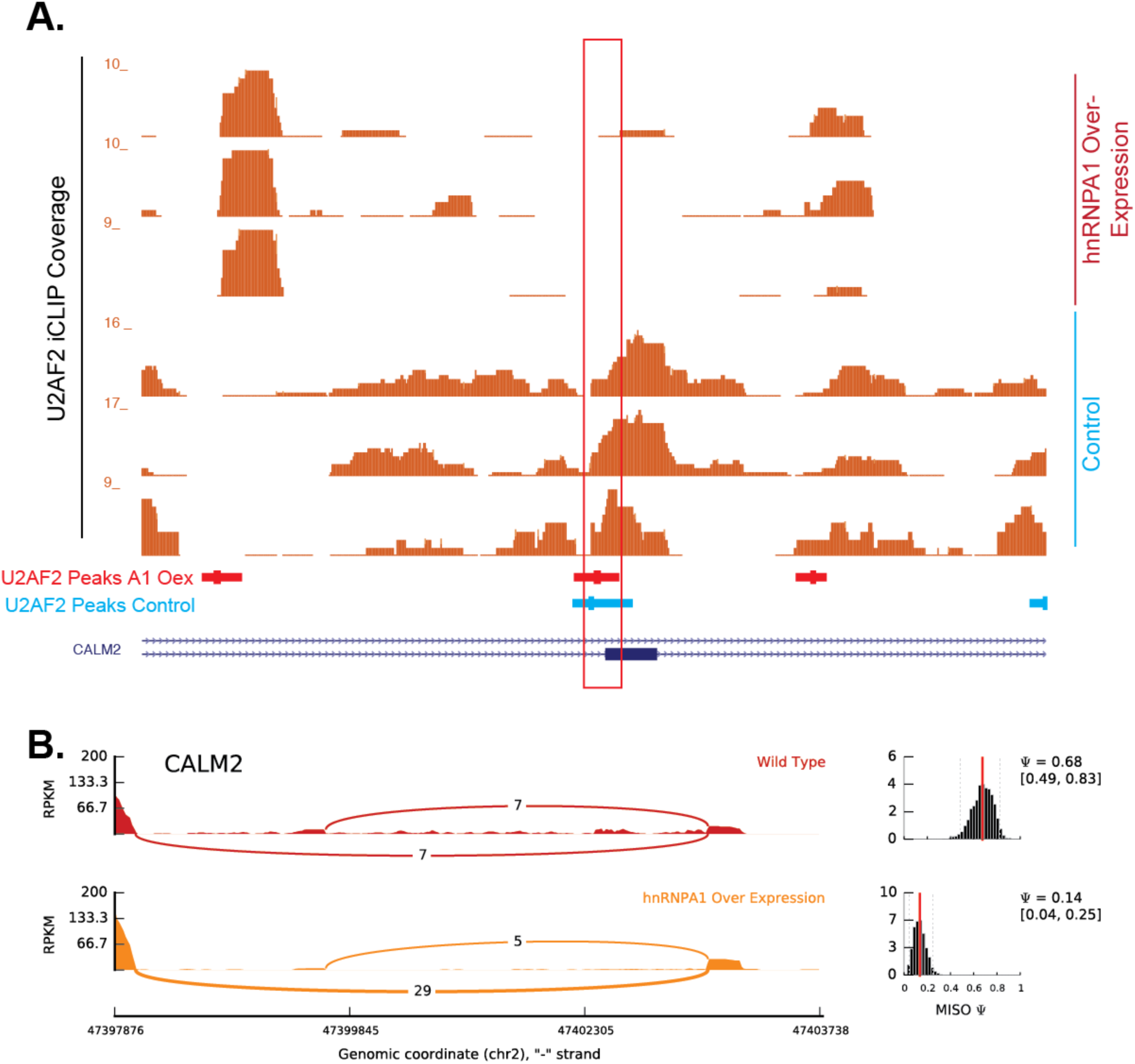
hnRNPA1-dependent modulation of U2AF2 crosslinking and alternative splicing of CALM2 locus. **(A)** UCSC genome browser of CALM2 alternative exon and CLIP read coverage data for U2AF2 under control (blue text) and hnRNPA1 overexpression (red text). Red box highlights region in which U2AF2 is seen to redistribute between two conditions. **(B)** Sashimi plots showing read and junction coverage for CALM2 gene under wildtype (red) and hnRNPA1 over-expression (orange) conditions. PSI (percent splicing index) values are provided for each condition reflecting inclusion level of exon shown in genome browser (A) image.

**Fig S8.**
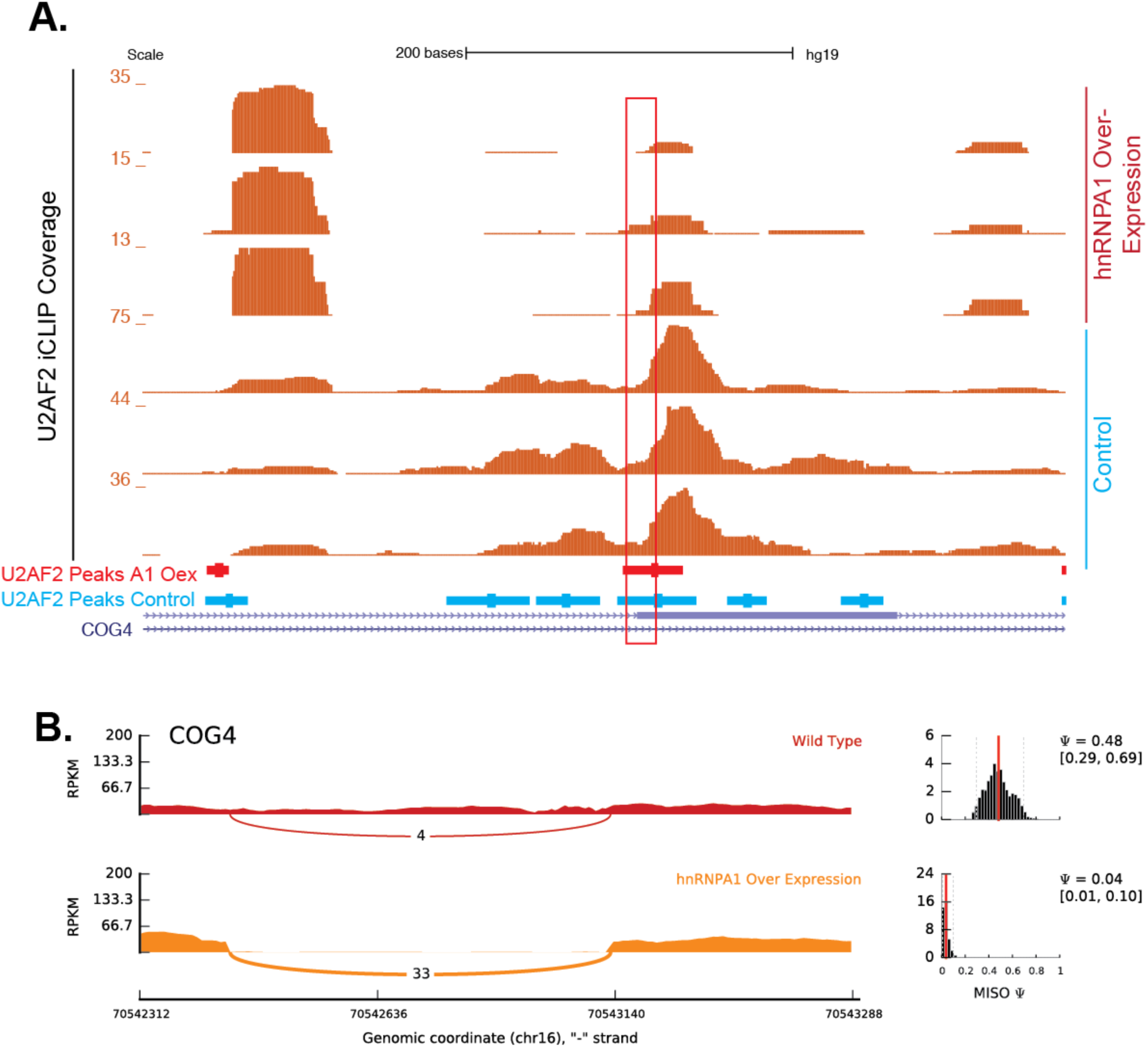
hnRNPA1-dependent modulation of U2AF2 crosslinking and alternative splicing of COG4 locus. **(A)** UCSC genome browser of COG4 alternative exon and CLIP read coverage data for U2AF2 under control (blue text) and hnRNPA1 overexpression (red text). Red box highlights region in which U2AF2 is seen to redistribute between two conditions. **(B)** Sashimi plots showing read and junction coverage for COG4 gene under wildtype (red) and hnRNPA1 over-expression (orange) conditions. PSI (percent splicing index) values are provided for each condition reflecting inclusion level of exon shown in genome browser (A) image.

**Fig S9.**
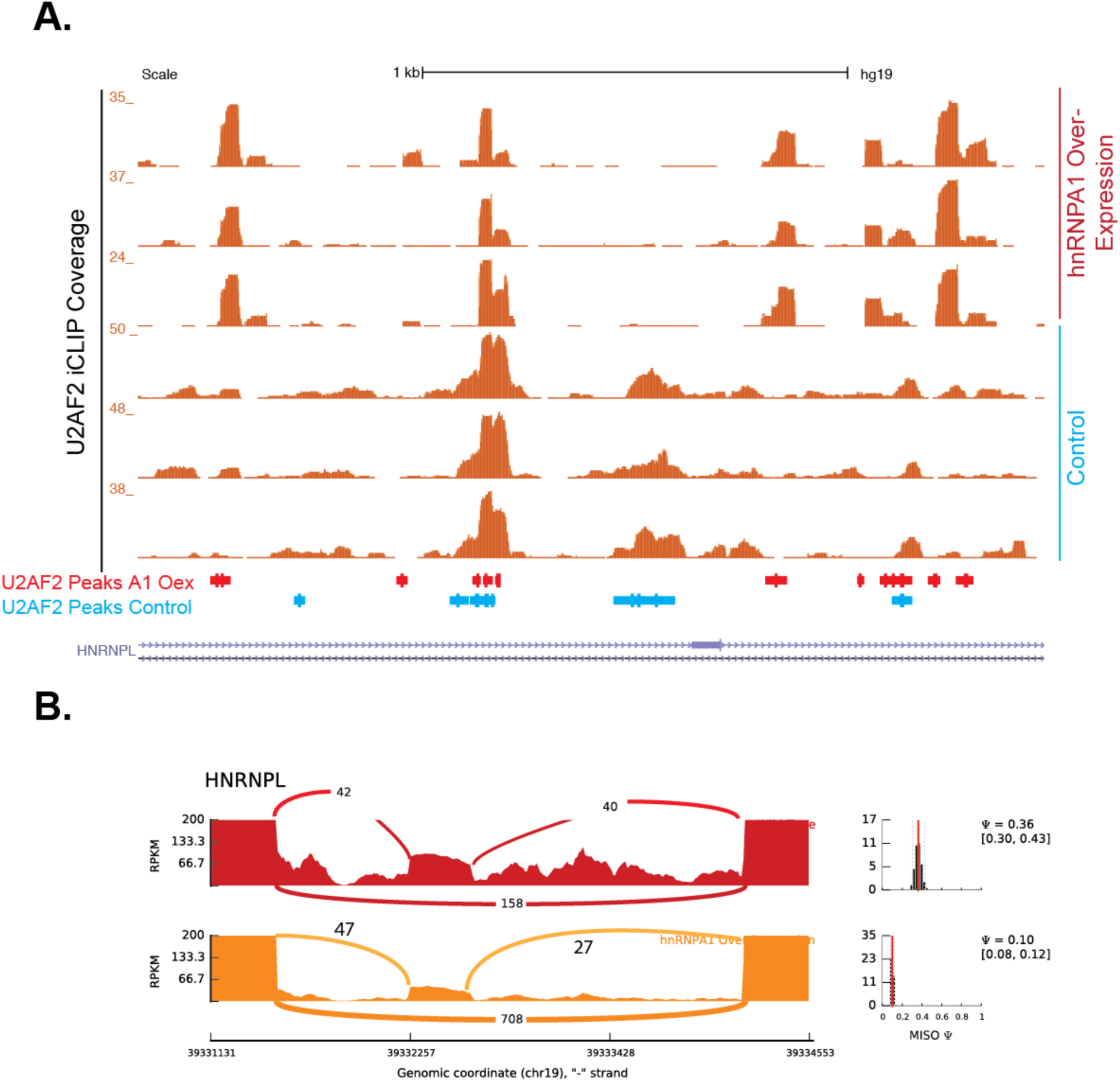
hnRNPA1-dependent modulation of U2AF2 crosslinking and alternative splicing of hnRNP L locus. **(A)** UCSC genome browser of COG4 alternative exon and CLIP read coverage data for U2AF2 under control (blue text) and hnRNPA1 overexpression (red text). Red box highlights region in which U2AF2 is seen to redistribute between two conditions. **(B)** Sashimi plots showing read and junction coverage for COG4 gene under wildtype (red) and hnRNPA1 over-expression (orange) conditions. PSI (percent splicing index) values are provided for each condition reflecting inclusion level of exon shown in genome browser (A) image.

**Fig S10.**
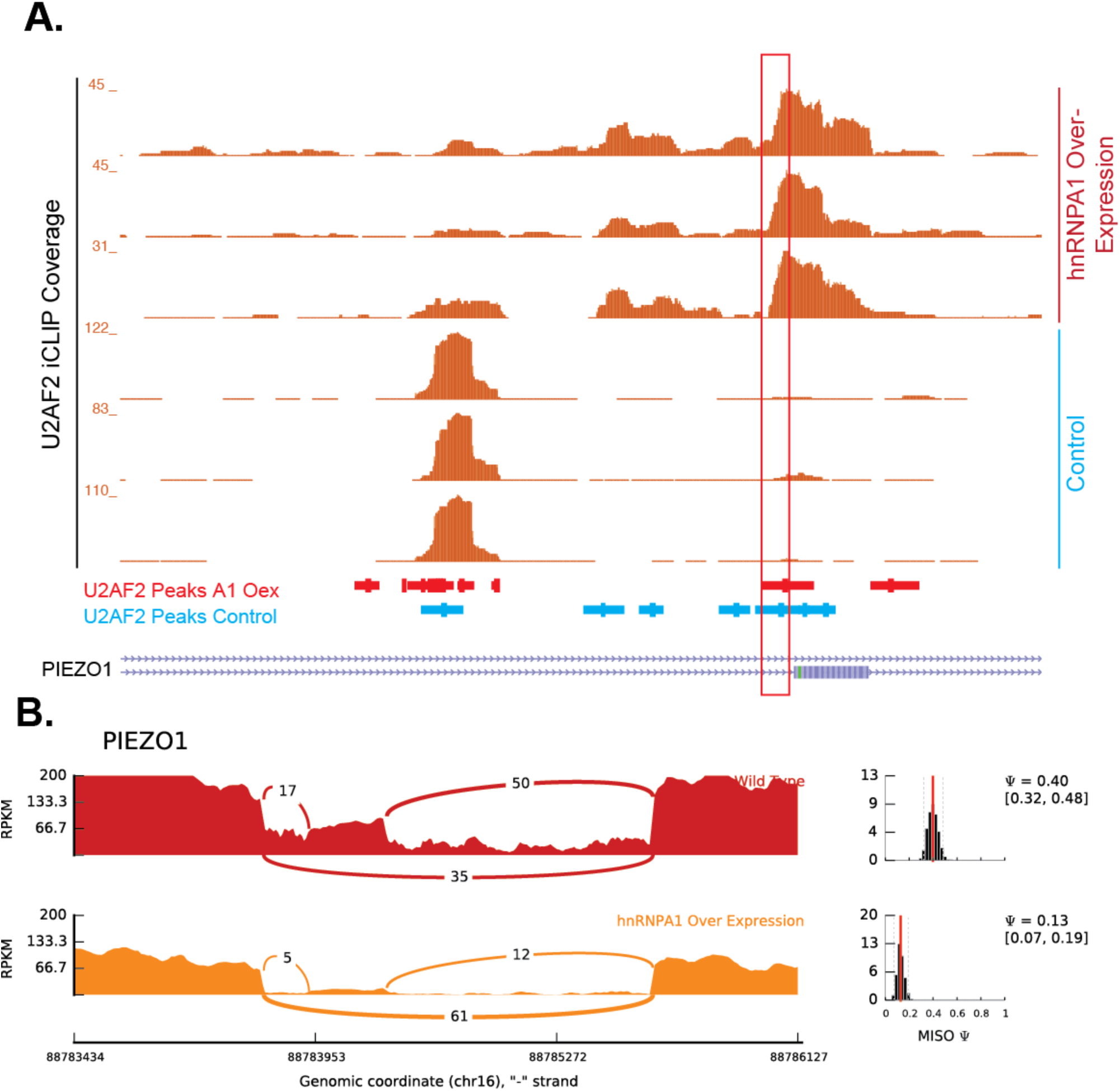
hnRNPA1-dependent modulation of U2AF2 crosslinking and alternative splicing of PIEZO1 locus. **(A)** UCSC genome browser of PIEZO1 alternative exon and CLIP read coverage data for U2AF2 under control (blue text) and hnRNPA1 overexpression (red text). Red box highlights region in which U2AF2 is seen to redistribute between two conditions. **(B)** Sashimi plots showing read and junction coverage for PIEZO1 gene under wildtype (red) and hnRNPA1 over-expression (orange) conditions. PSI (percent splicing index) values are provided for each condition reflecting inclusion level of exon shown in genome browser (A) image.

**Fig S11.**
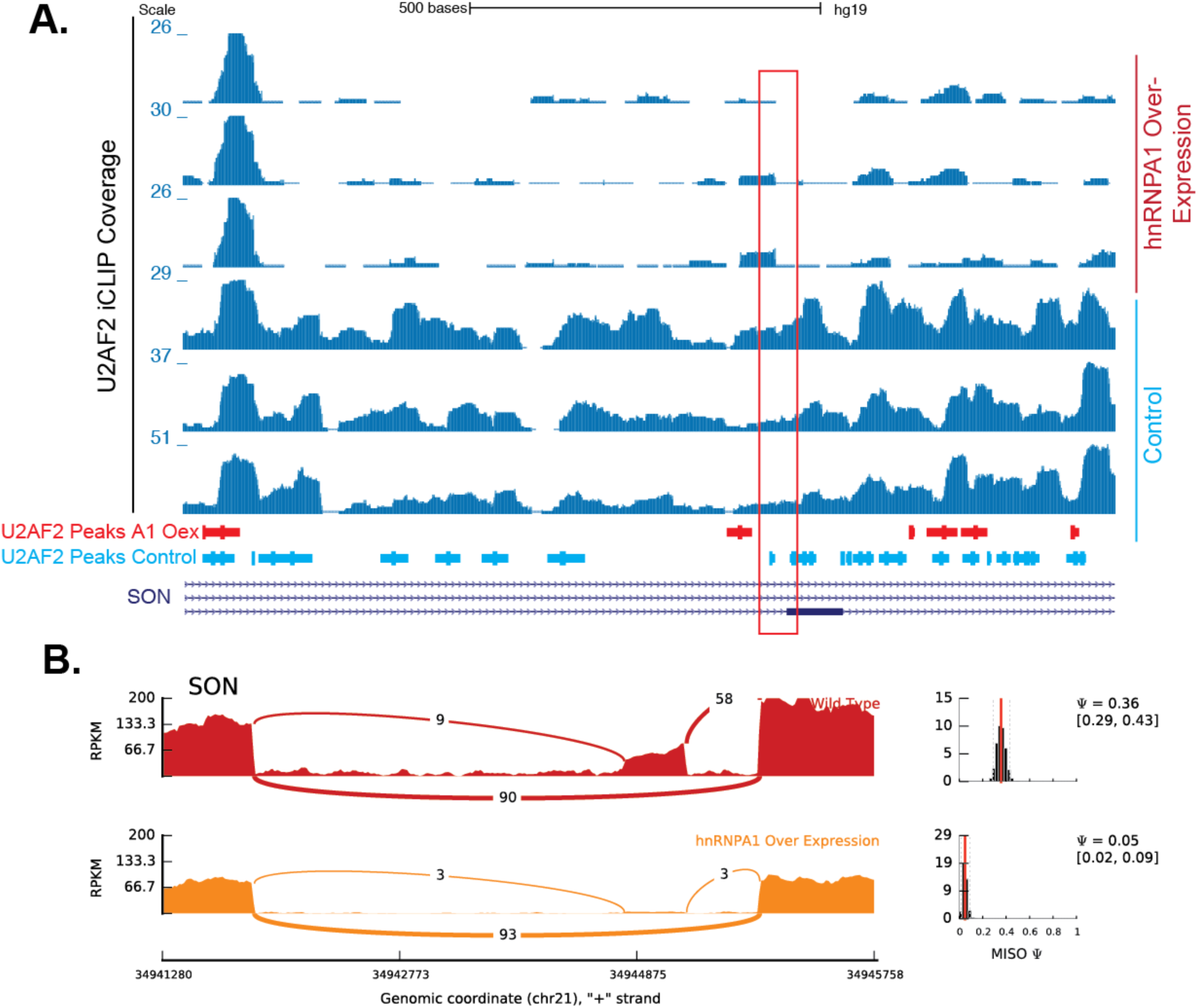
hnRNPA1-dependent modulation of U2AF2 crosslinking and alternative splicing of SON locus. **(A)** UCSC genome browser of SON alternative exon and CLIP read coverage data for U2AF2 under control (blue text) and hnRNPA1 overexpression (red text). Red boxes highlight region in which U2AF2 is seen to redistribute between two conditions. **(B)** Sashimi plots showing read and junction coverage for SON gene under wildtype (red) and hnRNPA1 over-expression (orange) conditions. PSI (percent splicing index) values are provided for each condition reflecting inclusion level of exon shown in genome browser (A) image.

**Fig S12.**
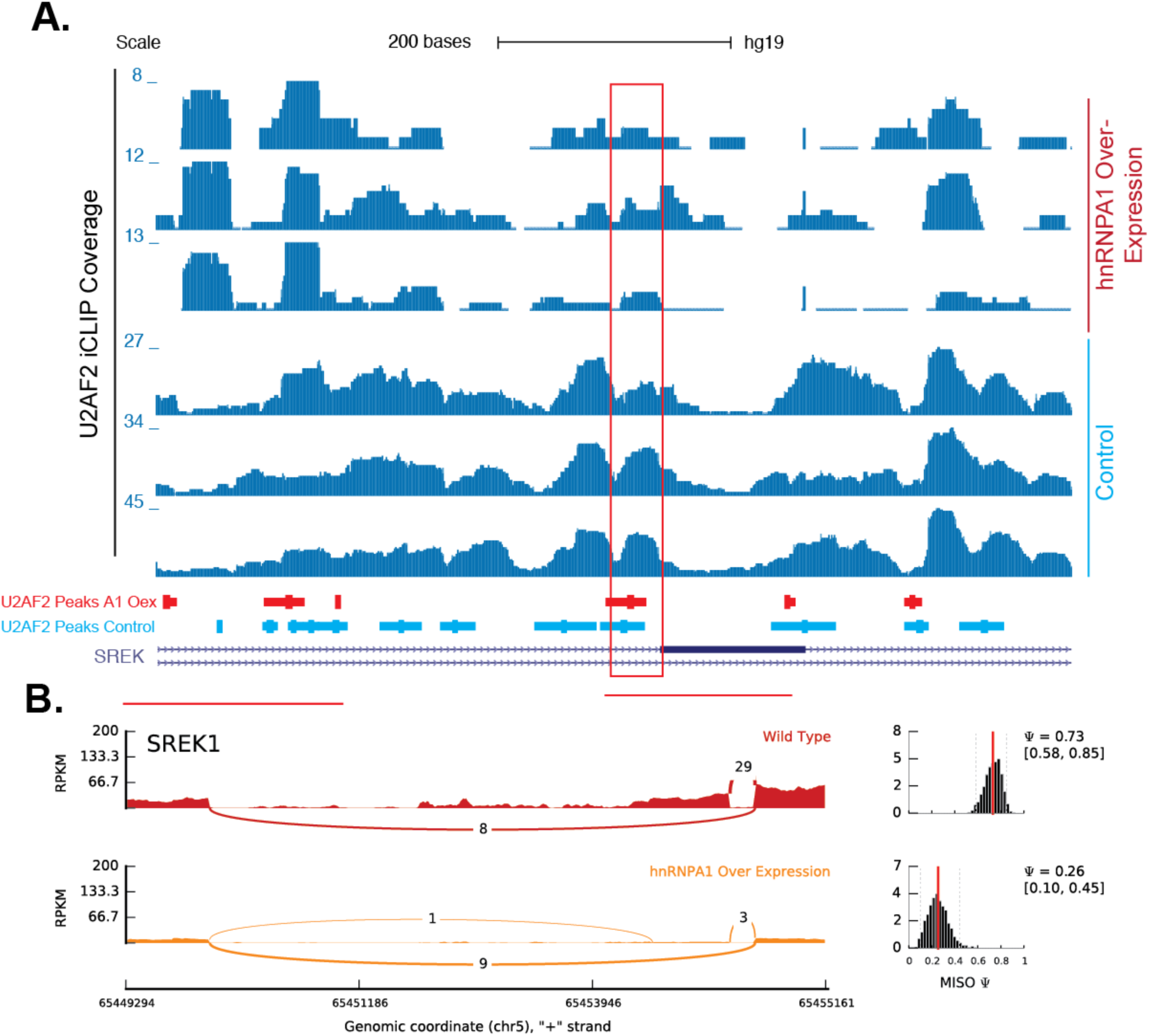
hnRNPA1-dependent modulation of U2AF2 crosslinking and alternative splicing of SREK1 locus. **(A)** UCSC genome browser of SREK1 alternative exon and CLIP read coverage data for U2AF2 under control (blue text) and hnRNPA1 overexpression (red text). Red boxes highlight region in which U2AF2 is seen to redistribute between two conditions. **(B)** Sashimi plots showing read and junction coverage for SREK1 gene under wildtype (red) and hnRNPA1 over-expression (orange) conditions. PSI (percent splicing index) values are provided for each condition reflecting inclusion level of exon shown in genome browser (A) image.

**Fig S13.**
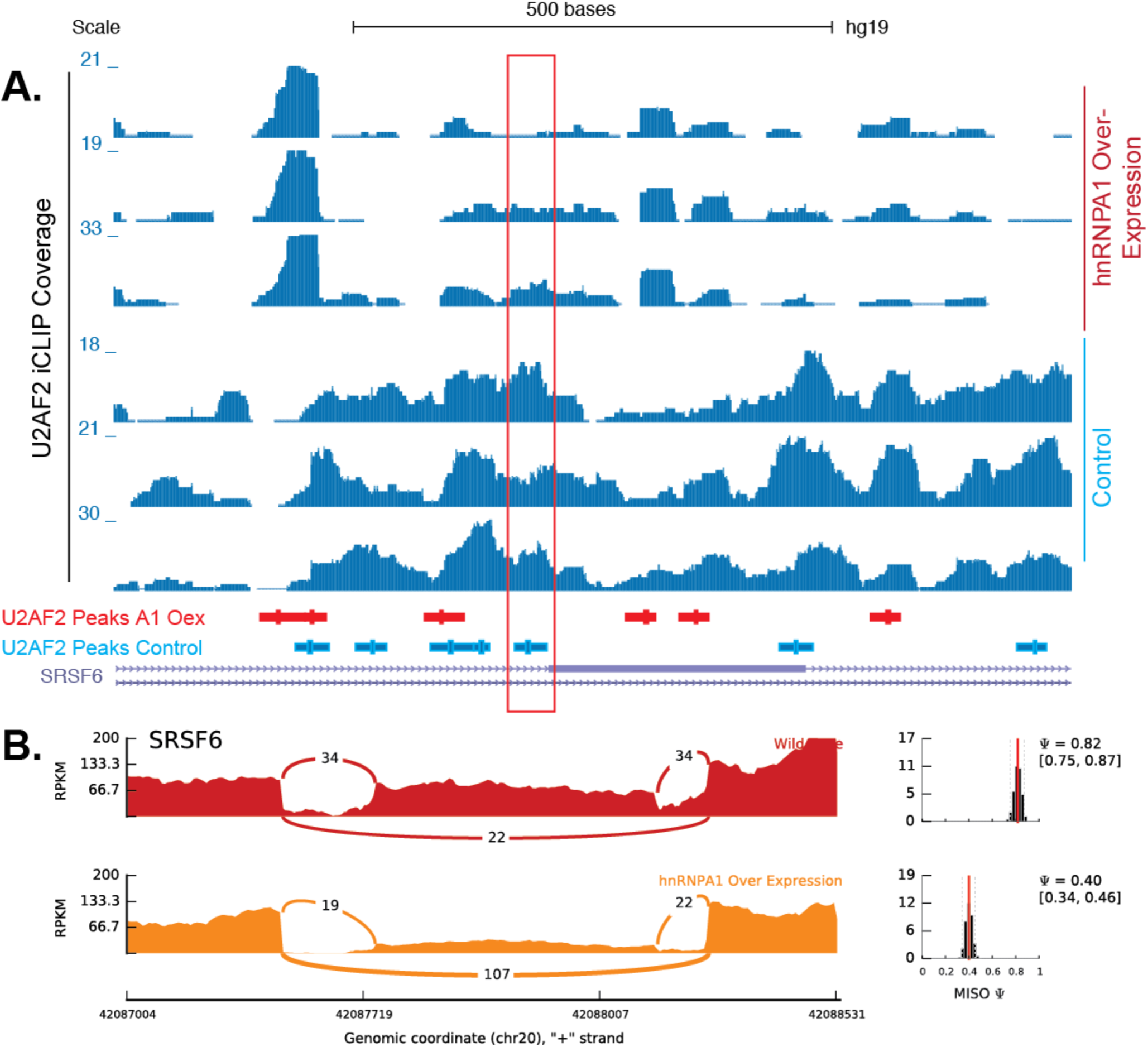
hnRNPA1-dependent modulation of U2AF2 crosslinking and alternative splicing of SRSF6 locus. **(A)** UCSC genome browser of SRSF6 alternative exon and CLIP read coverage data for U2AF2 under control (blue text) and hnRNPA1 overexpression (red text). **(B)** Sashimi plots showing read and junction coverage for SRSF6 gene under wildtype (red) and hnRNPA1 over-expression (orange) conditions. PSI (percent splicing index) values are provided for each condition reflecting inclusion level of exon shown in genome browser (A) image.

**Fig S14.**
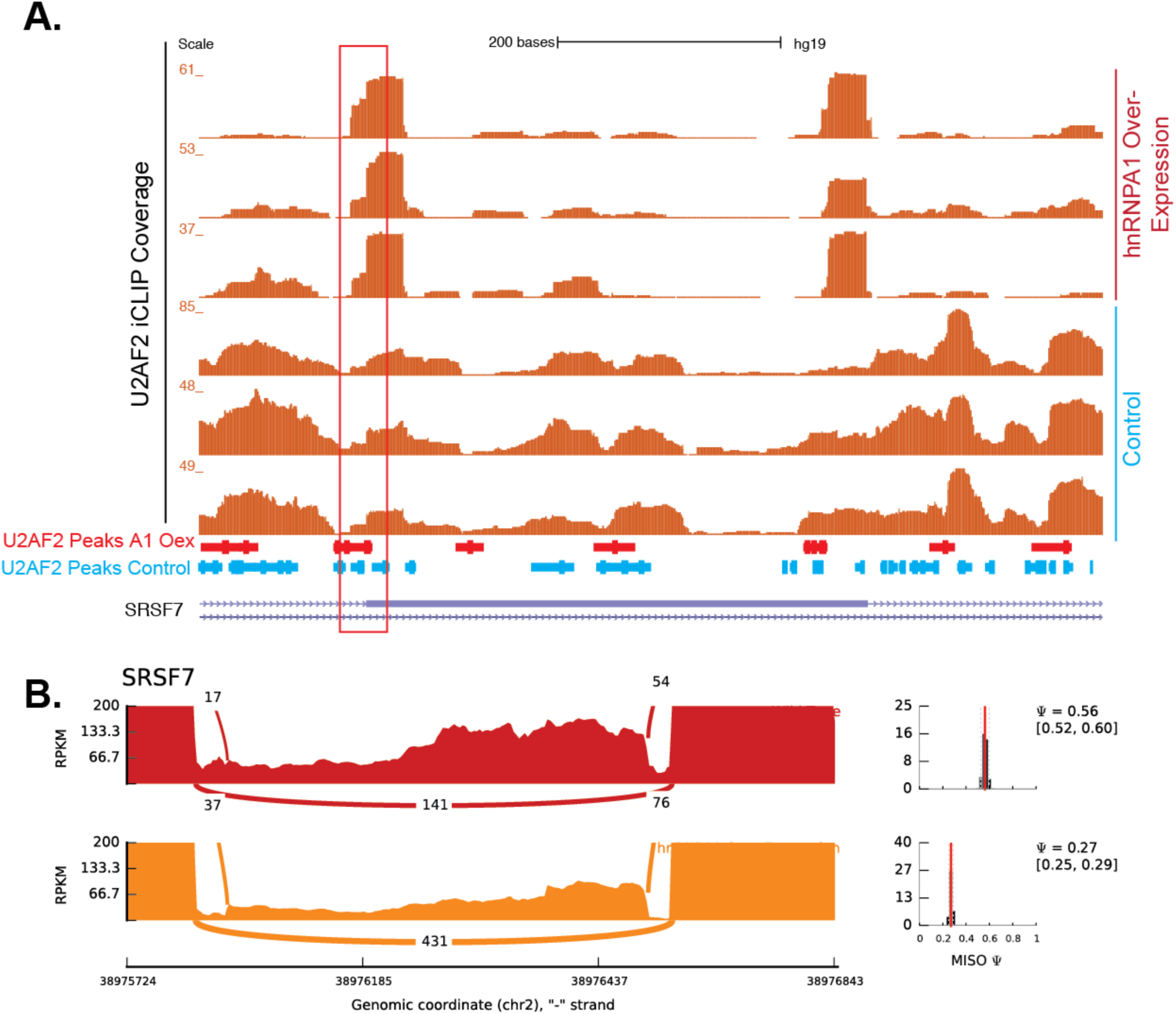
hnRNPA1-dependent modulation of U2AF2 crosslinking and alternative splicing of SRSF7 locus. **(A)** UCSC genome browser of SRSF7 alternative exon and CLIP read coverage data for U2AF2 under control (blue text) and hnRNPA1 overexpression (red text). **(B)** Sashimi plots showing read and junction coverage for SRSF7 gene under wildtype (red) and hnRNPA1 over-expression (orange) conditions. PSI (percent splicing index) values are provided for each condition reflecting inclusion level of exon shown in genome browser (A) image.

**Fig S15.**
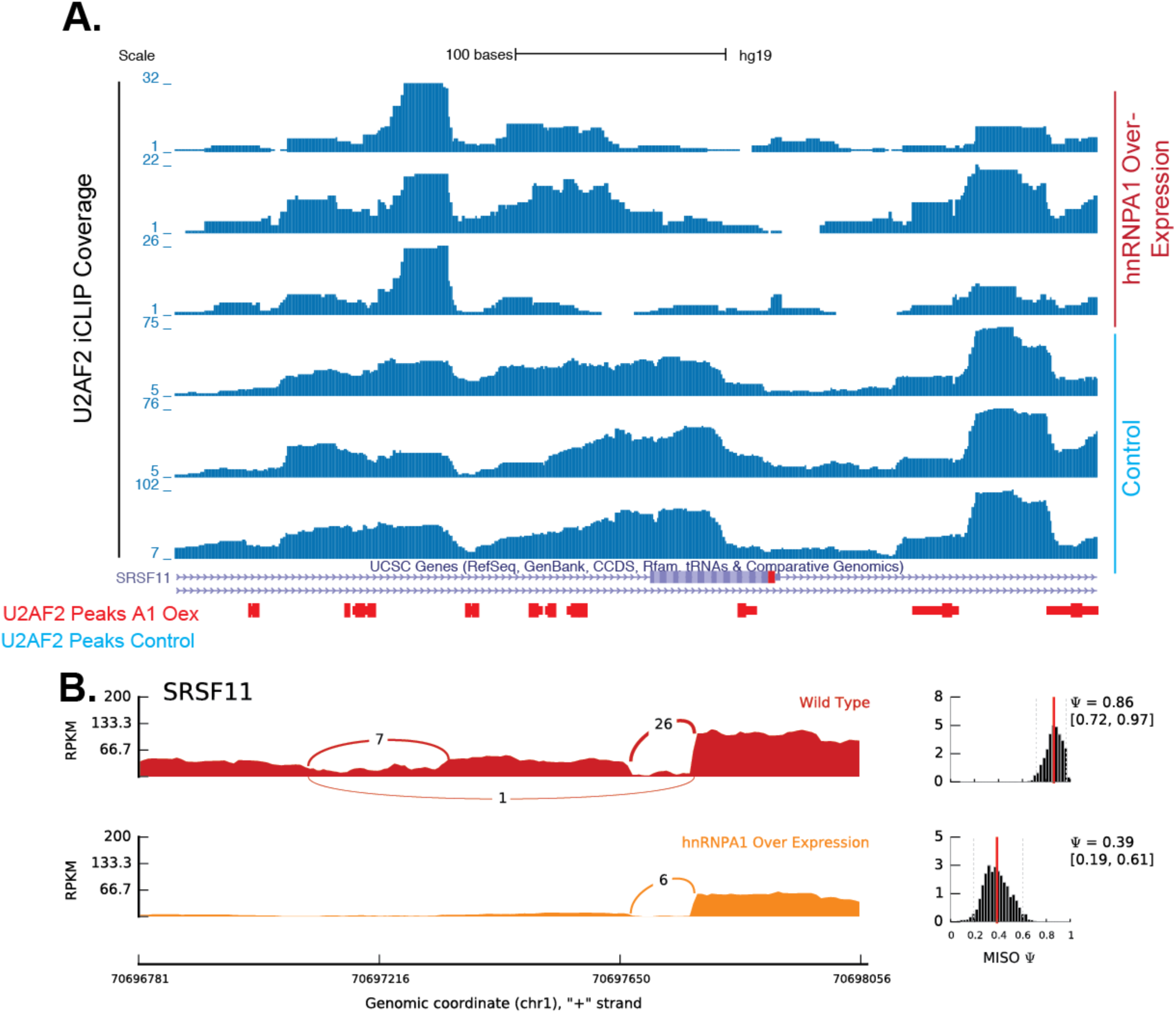
hnRNPA1-dependent modulation of U2AF2 crosslinking and alternative splicing of SRSF11 locus. **(A)** UCSC genome browser of SRSF11 alternative exon and CLIP read coverage data for U2AF2 under control (blue text) and hnRNPA1 overexpression (red text). **(B)** Sashimi plots showing read and junction coverage for SRSF11 gene under wildtype (red) and hnRNPA1 over-expression (orange) conditions. PSI (percent splicing index) values are provided for each condition reflecting inclusion level of exon shown in genome browser (A) image.

**Fig S16.**
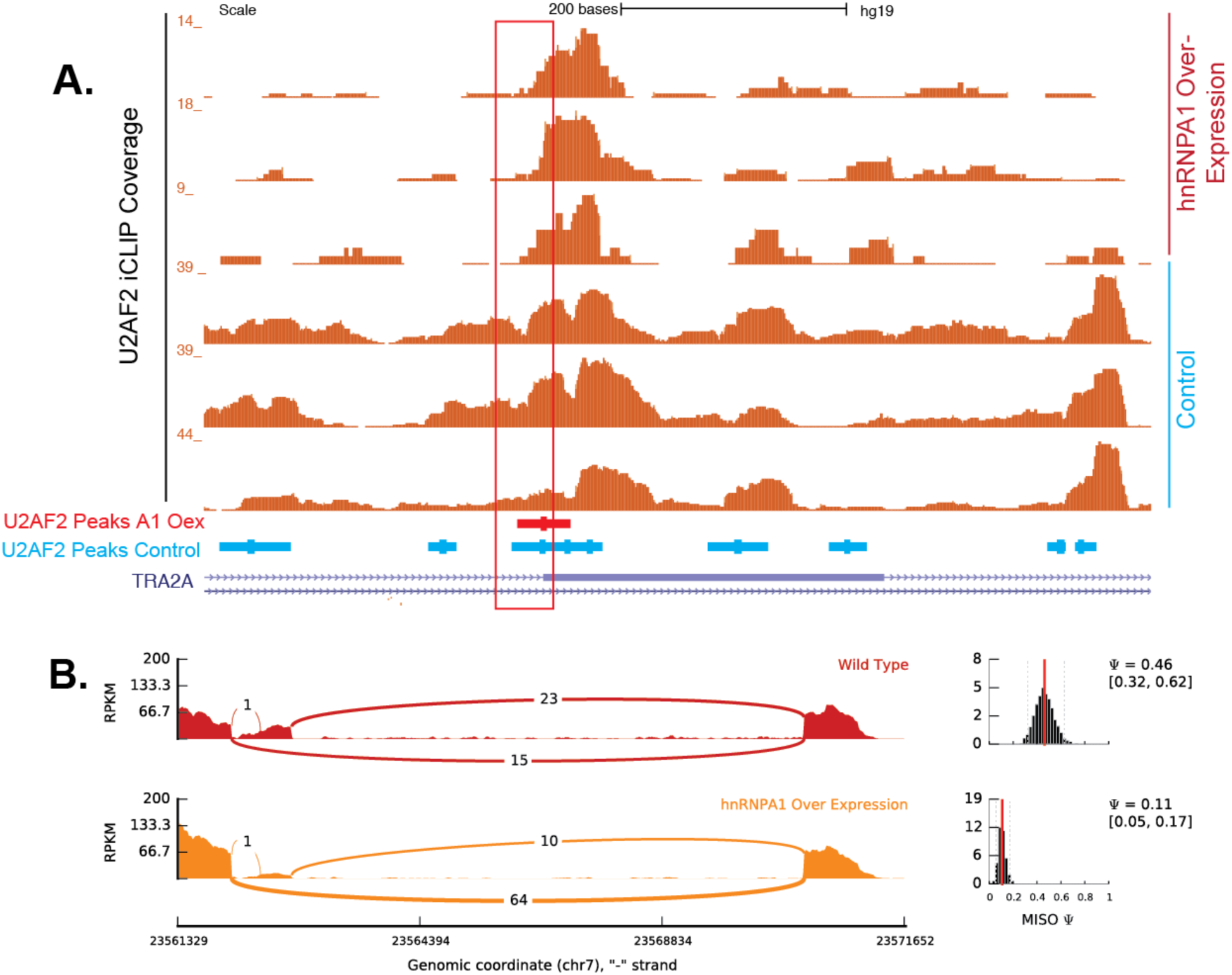
hnRNPA1-dependent modulation of U2AF2 crosslinking and alternative splicing of TRA2A locus. **(A)** UCSC genome browser of TRA2A alternative exon and CLIP read coverage data for U2AF2 under control (blue text) and hnRNPA1 overexpression (red text). **(B)** Sashimi plots showing read and junction coverage for TRA2A gene under wildtype (red) and hnRNPA1 over-expression (orange) conditions. PSI (percent splicing index) values are provided for each condition reflecting inclusion level of exon shown in genome browser (A) image.

**Fig S17.**
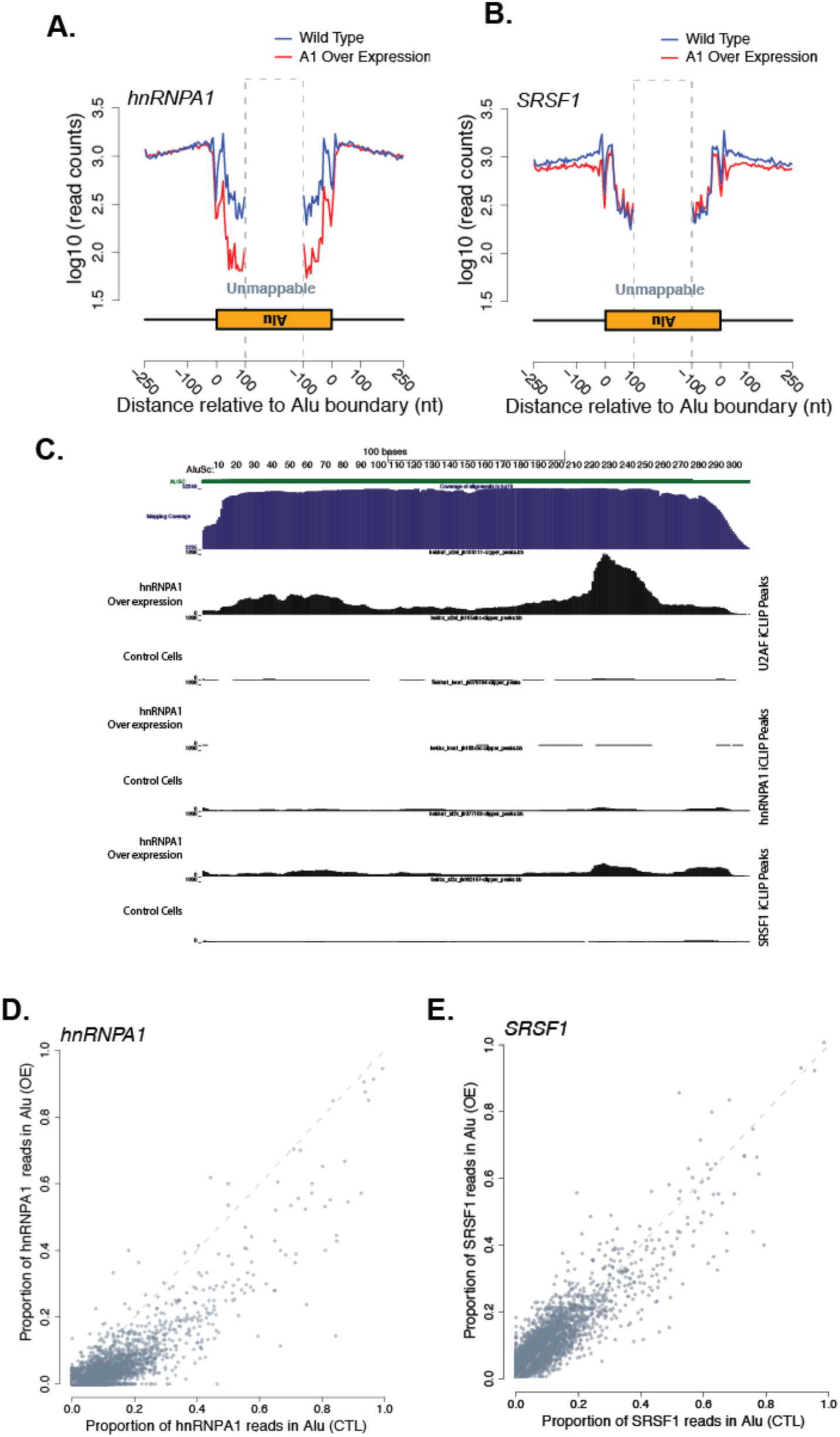
Global hnRNP A1 and SRSF1 crosslinking near antisense *Alu* elements under control and hnRNPA1 overexpression conditions. **(A,B)** Aggregated read counts on Alu elements and nearby regions for hnRNPA1 (A) and SRSF1 (B). Blue represents wild-type binding of the given RNA binding protein and red represents hnRNP A1 overexpression of the log10 number of iCLIP read counts across all antisense-Alu elements. **(C)** Distribution of aggregated U2AF2 (top), hnRNPA1 (middle), and SRSF1 (bottom) iCLIP read counts on Alu subtype *AluSc* under control and hnRNPA1 overexpression conditions. **(D,E)** Effects of hnRNP A1 overexpression on the proportion of hnRNP A1 and SRSF1 crosslinking sites in Alu RNA elements. Scatter plot of all human cassette exons measuring the proportion of hnRNP A1 and SRSF1 iCLIP crosslinking sites found within Alu elements relative to the total number of crosslinks observed throughout the alterative event.

## Supplemental Text

### Link UCSC Genome Browser session with RNA-seq coverage, iCLIP coverage and peaks

http://genome.ucsc.edu/cgi-bin/hgTracks?hgS_doOtherUser=submit&hgS_otherUserName=jeremyrsanford&hgS_otherUserSessionName=Howard%20et%20al

### iCLIP Coverage track descriptions plus strand

hekha1_hna1_jh104a ih1 rdcb1 1p cov = “hnRNPA1 iCLIP from hnRNPA1 overexpression cells” replicate 1

hekha1_hna1_jh104b ih1 rdcb1 1p cov= “hnRNPA1 iCLIP from hnRNPA1 overexpression cells” replicate 2

hektrx_hna1_jh103b ih1 rdcb1 1p cov= “hnRNPA1 iCLIP from control cells” replicate 1

hektrx_hna1_jh103c ih1 rdcb1 1p cov = “hnRNPA1 iCLIP from control cells” replicate 2

hekha1_sf2x_jh108a ih1 rdcb1 1p cov =“SRSF1 iCLIP from hnRNPA1 overexpression cells” replicate 1

hekha1_sf2x_jh108b ih1 rdcb1 1p cov =“SRSF1 iCLIP from hnRNPA1 overexpression cells” replicate 2

hekha1_sf2x_jh108c ih1 rdcb1 1p cov =“SRSF1 iCLIP from hnRNPA1 overexpression cells” replicate 3

hektrx_sf2x_jh107a ih1 rdcb1 1p cov =“SRSF1 iCLIP from control cells” replicate 1

hektrx_sf2x_jh107b ih1 rdcb1 1p cov =“SRSF1 iCLIP from control cells” replicate 2

hektrx_sf2x_jh107c ih1 rdcb1 1p cov =“SRSF1 iCLIP from control cells” replicate 3

hekha1_u2af_jh106a ih1 rdcb1 1p cov =“U2AF2 iCLIP from hnRNPA1 overexpression cells” replicate 1

hekha1_u2af_jh106b ih1 rdcb1 1p cov =“U2AF2 iCLIP from hnRNPA1 overexpression cells” replicate 2

hekha1_u2af_jh106c ih1 rdcb1 1p cov =“U2AF2 iCLIP from hnRNPA1 overexpression cells” replicate 3

hektrx_u2af_jh105a ih1 rdcb1 1p cov =“U2AF2 iCLIP from control cells” replicate 1 hektrx_u2af_jh105b ih1 rdcb1 1p cov =“U2AF2 iCLIP from control cells” replicate 2 hektrx_u2af_jh105c ih1 rdcb1 1p cov =“U2AF2 iCLIP from control cells” replicate 3

### Minus strand

hekha1_hna1_jh104a ih1 rdcb1 1m cov = “hnRNPA1 iCLIP from hnRNPA1 overexpression cells” replicate 1

hekha1_hna1_jh104b ih1 rdcb1 1m cov= “hnRNPA1 iCLIP from hnRNPA1 overexpression cells” replicate 2

hektrx_hna1_jh103b ih1 rdcb1 1m cov= “hnRNPA1 iCLIP from control cells” replicate 1

hektrx_hna1_jh103c ih1 rdcb1 1m cov = “hnRNPA1 iCLIP from control cells” replicate 2

hekha1_sf2x_jh108a ih1 rdcb1 1m cov =“SRSF1 iCLIP from hnRNPA1 overexpression cells” replicate 1

hekha1_sf2x_jh108b ih1 rdcb1 1m cov =“SRSF1 iCLIP from hnRNPA1 overexpression cells” replicate 2

hekha1_sf2x_jh108c ih1 rdcb1 1m cov =“SRSF1 iCLIP from hnRNPA1 overexpression cells” replicate 3

hektrx_sf2x_jh107a ih1 rdcb1 1m cov =“SRSF1 iCLIP from control cells” replicate 1

hektrx_sf2x_jh107b ih1 rdcb1 1m cov =“SRSF1 iCLIP from control cells” replicate 2

hektrx_sf2x_jh107c ih1 rdcb1 1m cov =“SRSF1 iCLIP from control cells” replicate 3

hekha1_u2af_jh106a ih1 rdcb1 1m cov =“U2AF2 iCLIP from hnRNPA1 overexpression cells” replicate 1

hekha1_u2af_jh106b ih1 rdcb1 1m cov =“U2AF2 iCLIP from hnRNPA1 overexpression cells” replicate 2

hekha1_u2af_jh106c ih1 rdcb1 1m cov =“U2AF2 iCLIP from hnRNPA1 overexpression cells” replicate 3

hektrx_u2af_jh105a ih1 rdcb1 1m cov =“U2AF2 iCLIP from control cells” replicate 1

hektrx_u2af_jh105b ih1 rdcb1 1m cov =“U2AF2 iCLIP from control cells” replicate 2

hektrx_u2af_jh105c ih1 rdcb1 1m cov = “U2AF2 iCLIP from control cells” replicate 3

### CLIPper Peak Track descriptions

hekha1_hna1_jh078104 P2 clipper_pk = Peaks called from hnRNPA1 iCLIP from hnRNPA1 over-expression cells

hektrx_hna1_jh103xbc P2 clipper_pk = Peaks called from hnRNPA1 iCLIP from control cells hekha1_sf2x_jh077108 P2 clipper_pk = Peaks called from SRSF1 iCLIP from hnRNPA1 over-expression cells

hektrx_sf2x_jh082107 P2 clipper_pk = Peaks called from SRSF1 iCLIP from control cells hekha1_u2af_jh106111 P2 clipper_pk = Peaks called from U2AF2 iCLIP from hnRNPA1 over-expression cells

hektrx_u2af_jh105abc P2 clipper_pk = Peaks called from U2AF2 iCLIP from control cells

### RNA-Seq Coverage Tracks

hekha1_clip_ti_jh45 ih1 rd1 xpe 0b cov = RNA-seq from hnRNP over-expression cells replicate 1

hekha1_clip_ti_jh46 ih1 rd1 xpe 0b cov = RNA-seq from hnRNP over-expression cells replicate 2

hekha1_clip_tn_jh43 ih1 rd1 xpe 0b cov = RNA-seq from control cells replicate 1 hekha1_clip_tn_jh44 ih1 rd1 xpe 0b cov = RNA-seq from control cells replicate 2

